# Microbial interactions impact the growth response of *Clostridioides difficile* to antibiotics

**DOI:** 10.1101/2022.09.16.508345

**Authors:** Susan Hromada, Ophelia Venturelli

**Affiliations:** Department of Biochemistry, University of Wisconsin-Madison, Madison, WI, USA; Microbiology Doctoral Training Program, University of Wisconsin-Madison, WI, USA; Department of Bacteriology, University of Wisconsin-Madison, Madison, WI, USA; Department of Chemical and Biological Engineering, University of Wisconsin-Madison, Madison, WI, USA

## Abstract

In the human gut, the growth of *Clostridioides difficile* is impacted by a complex web of inter-species interactions with members of human gut microbiota. We investigate the contribution of inter-species interactions on the antibiotic response of *C. difficile* to clinically relevant antibiotics using bottom-up assembly of human gut communities. We discover two classes of microbial interactions that alter *C.* difficile’s antibiotic susceptibility: infrequent increases in tolerance at high antibiotic concentrations and frequent growth enhancements at low antibiotic concentrations. Based on genome-wide transcriptional profiling data, we demonstrate that metal sequestration due to hydrogen sulfide production by the prevalent gut species *Desulfovibrio piger* increases metronidazole tolerance of *C. difficile*. Competition with species that display higher sensitivity to the antibiotic than *C. difficile* leads to enhanced growth of *C. difficile* at low antibiotic concentrations. A dynamic computational model identifies the ecological design principles driving this effect. Our results provide a deeper understanding of ecological and molecular principles shaping *C. difficile*’s response to antibiotics, which could inform therapeutic interventions.

## INTRODUCTION

The bacterial pathogen *C. difficile* can infect the human gastrointestinal tract, an environment teeming with a dense microbiota. The gut microbiota has significant interactions with *C. difficile* which inhibit *C. difficile*‘s growth and ability to persist over time in the human gut, a phenomenon known as colonization resistance^1^. The key role of colonization resistance is illustrated by the increased risk of *C. difficile* infection after treatment with antibiotics that decimate the microbiota^2^ and by the efficacy of fecal microbiota transplants from healthy human donors to eliminate *C. difficile* infections^3^. Previous studies have provided a deeper understanding of interactions between gut microbiota and *C. difficile*. For example, inter-species interaction between individual gut microbes and *C. difficile* have been studied *in vitro* and analyses of human microbiome data have identified gut microbes whose presence or absence is associated with *C. difficile* infection^4–6^. Multiple mechanisms of interaction have been determined, such as inhibition of *C. difficile* germination by *Clostridium scindens* via production of secondary bile acids^6^ and promotion of *C. difficile* growth by *Bacteroides thetaiotaomicron* via succinate cross-feeding^7^. While much is known about how the microbiota impacts *C. difficile* growth, how the microbiota impacts *C. difficile* antibiotic susceptibility is largely unknown.

Understanding how the gut microbiota impacts *C. difficile* antibiotic susceptibility is important because antibiotics are the most widely used treatment for *C. difficile* infections. For example, vancomycin and metronidazole are two antibiotics used to treat *C. difficile* infections. Vancomycin is a glycopeptide that inhibits cell-wall synthesis whose activity is specific to gram-positive bacteria^8^. Metronidazole is a DNA-damaging agent that is effective against both gram-positive and gram-negative anaerobic bacteria. Metronidazole is a prodrug which is inactive until its nitro group is reduced to nitroso radicals in the cytoplasm of anaerobic bacteria^9^. The proposed mechanism of metronidazole reduction is due to the cofactors ferredoxin and/or flavodoxin in reactions catalyzed by multiple enzymes including reductases, hydrogenases and pyruvate ferredoxin/flavodoxin oxidoreductase (PFOR)^10–12^. Metronidazole, previously a recommended first-line treatment, is now only recommended in rare cases due to an observed decrease in its clinical effectiveness^13, 14^.

Similar to other pathogens, *C. difficile* antibiotic susceptibility has been studied using *in vitro* experiments of monoculture growth. However, monoculture experiments do not consider how interactions with resident community members can modify the antibiotic susceptibility of a pathogen. For example, monospecies antibiotic susceptibility of the pathogen *Pseudomonas aeruginosa* did not always correlate with the efficacy of treatment for polymicrobial infections^15^. If microbial interactions significantly increase the pathogen’s antibiotic tolerance, treatments based on monoculture susceptibility to antibiotics may not be effective in eradicating the pathogen. Alternatively, if communities reduce the pathogen’s antibiotic tolerance, the standard antibiotic dosage may exceed the dose needed to eradicate the pathogen, yielding unnecessary and avoidable disruption to the native microbiota. Understanding how constituent members of microbiota alter a pathogen’s susceptibility to an antibiotic could be used to guide improved antibiotic treatments.

Previous studies have shown that inter-species interactions can alter a given microbe’s response to antibiotics by increasing or decreasing susceptibility compared to monoculture^16^. One example is exposure protection, where susceptible microbes are protected from an antibiotic by species that degrade the antibiotic^17^. In addition, a previous study showed that an increase in antibiotic susceptibility occurred when the growth of a resistant microbe depended on cross-feeding with another organism that was susceptible to the antibiotic^18^.

While previous studies have identified specific types of inter-species interactions that impact antibiotic susceptibility, the prevalence of susceptibility-altering microbial interactions across different microbial communities is not well understood. In a human urinary tract infection community, around a third of total interactions between species were estimated to yield a change in the susceptibility to two different antibiotics based on spent media experiments^19^. Studies of a fruit fly microbiome, a multispecies wound infection biofilm, and a multispecies brewery biofilm each identified a change in antibiotic susceptibility of a given species in the community compared to monospecies^20–22^. In contrast, no significant changes in antibiotic susceptibility were observed for 15 characterized species in a community of human gut microbes or an *E. coli* pathogen introduced into a porcine microbiome^23, 24^. Based on this observed variation in the contribution of inter-species interactions to antibiotic susceptibility across different systems, it is not known whether microbial interactions can impact *C. difficile*’s antibiotic susceptibility.

We use a diverse human gut community^4^ to study the impact of microbial interactions on *C. difficile’s* antibiotic susceptibility. Using the clinically relevant antibiotics metronidazole and vancomycin, we identify a general class of competitive inter-species interactions that lead to enhanced growth of *C. difficile* in low antibiotic concentrations below the minimum inhibitory concentration (MIC) compared to the absence of antibiotic. This mechanism stems from a reduction in the strength of biotic inhibition of *C. difficile* by antibiotic-sensitive competing species in the presence of low antibiotic concentrations. In the presence of high antibiotic concentrations, we demonstrate that the gut species *Desulfovibrio piger* can substantially enhance *C. difficile*’s tolerance to metronidazole. *D. piger* induces a metal starvation transcriptional response in *C. difficile*, which in turn yields a down-regulation of metal-containing enzymes hypothesized to reduce metronidazole to its active form, yielding a lower concentration of active antibiotic. Corroborating this notion, supplementation of *D. piger*’s supernatant with a combination of metals eliminates the observed increase in metronidazole tolerance. In sum, a deeper understanding of the role of inter-species interactions in shaping *C. difficile* ‘s response to antibiotics could inform therapeutic interventions.

## RESULTS

### C. difficile tolerance to metronidazole and vancomycin is modified by a subset of gut microbes in pairwise communities

*C. difficile* infections occur in the context of complex resident gut communities. However, antibiotic treatments to eliminate *C. difficile* infection are designed based on the susceptibility of *C. difficile* in monoculture, which neglect the role of inter-species interactions in shaping antibiotic susceptibility. Understanding how the gut microbiota alters *C. difficile* antibiotic susceptibility could inform the treatment of *C. difficile* by antibiotics. To investigate this question, we evaluated *C. difficile’s* response to antibiotics in the presence of synthetic communities of gut microbes (**Figure 1A**). The 13-member human gut community was designed to span the phylogenetic diversity of the human gut microbiome and the interactions between these gut microbes and *C. difficile* have been deciphered in the absence of antibiotics^4^ (**Figure 1B**). We selected the antibiotics metronidazole and vancomycin for their clinical relevance in the treatment of *C. difficile* infections and for the differences in their activity spectra. Metronidazole has broad spectrum activity against the 13 gut microbes, whereas the activity of vancomycin is specific for gram-positive species. Since *C. difficile* can encounter both low and high antibiotic concentrations in the human gut^25^, we characterized *C. difficile*’s minimum inhibitory concentration (MIC) and its abundance at sub-inhibitory concentrations across a range of ecological contexts (**Figure 1C**).

**Figure 1:**
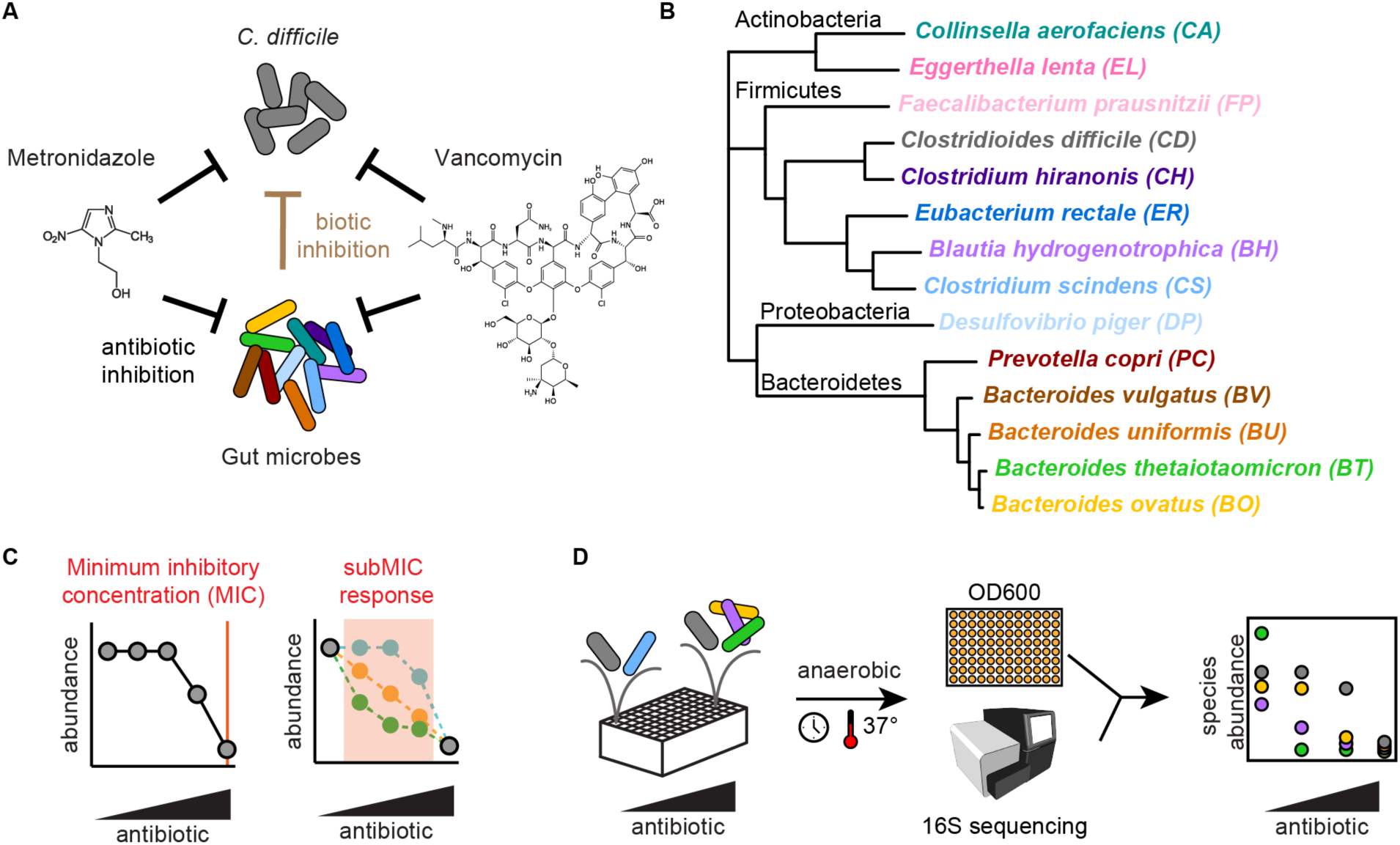
*C. difficile* antibiotic response in multispecies gut microbe communities. **(a)** Schematic of factors affecting *C. difficile* growth in synthetic gut communities in the presence of antibiotics metronidazole or vancomycin. Antibiotic inhibition (black arrows) by the DNA-damaging agent metronidazole and the cell wall inhibitor vancomycin inhibit *C. difficile* and gut microbes. Biotic inhibition by gut microbes (brown arrow) inhibits *C. difficile.* **(b)** Phylogenetic tree of 14-membered synthetic community of human gut microbes, spanning 4 major phyla of gut microbiota, and pathogen *C. difficile*. Black text indicates phylum name. Colored text indicates species name. Phylogeny based on concatenated alignment of 37 marker genes^26^. **(c)** Schematic of two aspects of the response of a microbial population to antibiotics: the minimum inhibitory concentration (MIC) and the sub-minimum inhibitory concentration (subMIC) response. **(d)** Schematic of methods for determining antibiotic susceptibility of multispecies communities. Communities are incubated anaerobically in microtiter plates. Absolute abundance is determined by multiplying optical density (OD600) by relative abundance from 16S rRNA gene sequencing.

To determine the antibiotic susceptibility of *C. difficile* in communities, we used a broth dilution method based on the clinical method detailed by the Clinical and Laboratory Standards Institute^27^ (Methods). We inoculated communities in liquid culture containing a twofold dilution series of antibiotics. After incubation, we determined the absolute abundance of each species in the communities at a fixed timepoint (48 hours) by multiplying the community optical density at 600 nm (OD600) by its relative abundance determined via 16S rRNA gene sequencing (**Figure 1D**). The MIC for each species in the community was determined as the lowest antibiotic concentration where growth at 48 hours was below a threshold (Methods).

Using this method, we determined the susceptibility of each species in monoculture (**Figure 2A**) and in each pairwise community containing *C. difficile* (**Figure 2BC**). Consistent with the mechanism of the antibiotics, all gut species were susceptible to metronidazole in the range of antibiotics tested, whereas gram-negative species (Bacteroidetes and Proteobacteria phyla, **Figure 1D**) were not susceptible to vancomycin (**Figure 2A**, **Supplementary Figure 1AB**).

**Figure 2:**
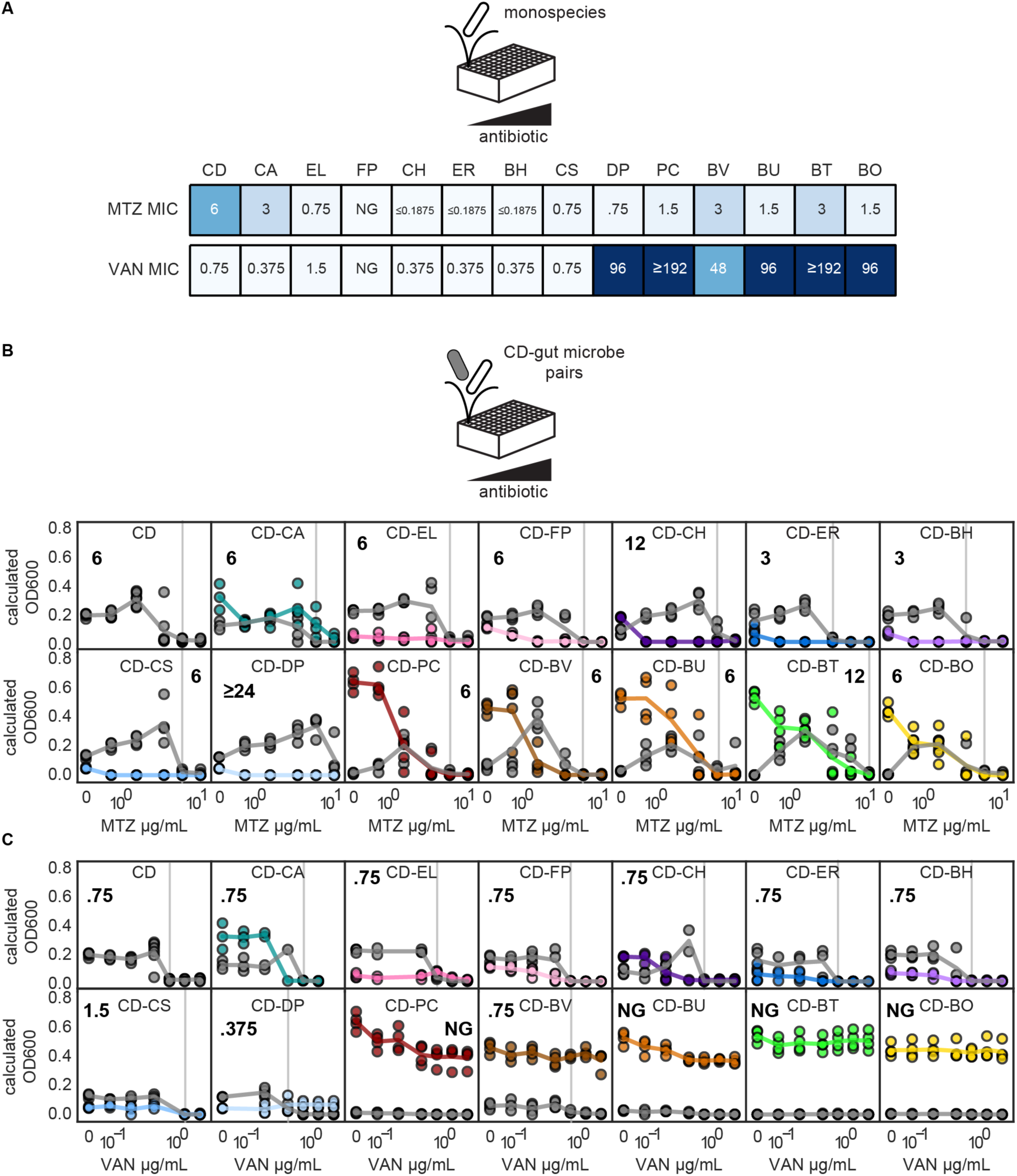
*C. difficile* response to metronidazole and vancomycin is modified by a subset of gut microbes in pairwise communities. **(a)** Heatmaps of minimum inhibitory concentration (MIC) of monocultures in the presence of metronidazole (MTZ) or vancomycin (VAN). Species indicated by two letter species code (Fig. 1B). **(b) (c)** Line plots of species absolute abundance at 48 hours as a function of antibiotic concentration for *C. difficile* monospecies or *C. difficile* in pairwise communities with another member of the human gut community. Each x-axis is semi-log scale. The values on the y-axis are calculated OD600 (OD600 multiplied by relative abundance from 16S rRNA gene sequencing). Data points indicate individual biological replicates. Lines indicate the average of n=1 to n=8 biological replicates. Vertical gray line and bold number indicate MIC of *C. difficile* in each condition. Color indicates species (Fig. 1B).

Substantial changes in *C. difficile*’s MIC in the presence of other gut microbes could lead to ineffective treatment if *C. difficile* is protected from the action of the antibiotic. Alternatively, if *C. difficile* tolerance is decreased in the community context, the concentration of the antibiotic is therefore higher than needed, leading to unnecessary disruption of gut microbiota. In most pairwise communities, the MIC of *C. difficile* was unchanged compared to monoculture. The MIC of *C. difficile* was unchanged in 15 pairs and displayed a moderate difference (2-fold) in 6 pairs compared to monospecies. However, *C. difficile* was significantly protected from metronidazole in co-culture with *D. piger* with an increase in MIC of ≥4-fold (**Figure 2B**). In sum, the MIC of *C. difficile* was infrequently impacted by pairwise inter-species interactions in the human gut community.

### Enhancement of C. difficile abundance in pairwise communities in the presence of low antibiotic concentrations

In the gut, pathogens such as *C. difficile* can encounter antibiotic concentrations lower than the MIC at the beginning and end of dosing regimens and between daily dosages^25^. Further, if the pathogen has gained resistance to the antibiotic, the entire dose regimen could be sub-inhibitory. Therefore, we investigated the role of microbial interactions on *C. difficile’*s response to sub-inhibitory antibiotic concentrations (concentrations lower than *C. difficile*’s MIC or subMICs).

In many pairwise communities, *C. difficile’*s abundance was similar or reduced in the subMIC range than in the absence of antibiotic (**Figure 2BC**). However, in a subset of pairwise communities, *C. difficile*’s abundance in the subMIC range was enhanced compared to the absence of antibiotic (**Figure 2BC**). For example, *C. difficile* monospecies abundance was similar in the presence of 0.75 µg/mL metronidazole and the absence of antibiotic. However, in the presence of *B. thetaiotaomicron*, the abundance of *C. difficile* was significantly higher in the presence of 0.75 µg/mL metronidazole than in the absence of antibiotic (**Figure 2B**). To provide further insights, we computed the subMIC fold change at each subMIC, defined as the species absolute abundance at the given subMIC divided by the species absolute abundance in the absence of antibiotic (**Figure 3A**). Using this metric, *C. difficile*’s growth was enhanced for at least one subMIC in one pairwise community in the presence of vancomycin and 11 pairwise communities in the presence of metronidazole (**Figure 3B**, **Supplementary Figure 2AB**). In seven pairwise communities, the growth enhancement of *C. difficile* in the presence of metronidazole was significantly greater than the enhancement observed for the *C. difficile* monoculture in the presence of metronidazole (**Figure 3B**). In sum, our results illustrate that approximately half of the species altered *C. difficile*’s response to low concentrations of metronidazole.

**Figure 3:**
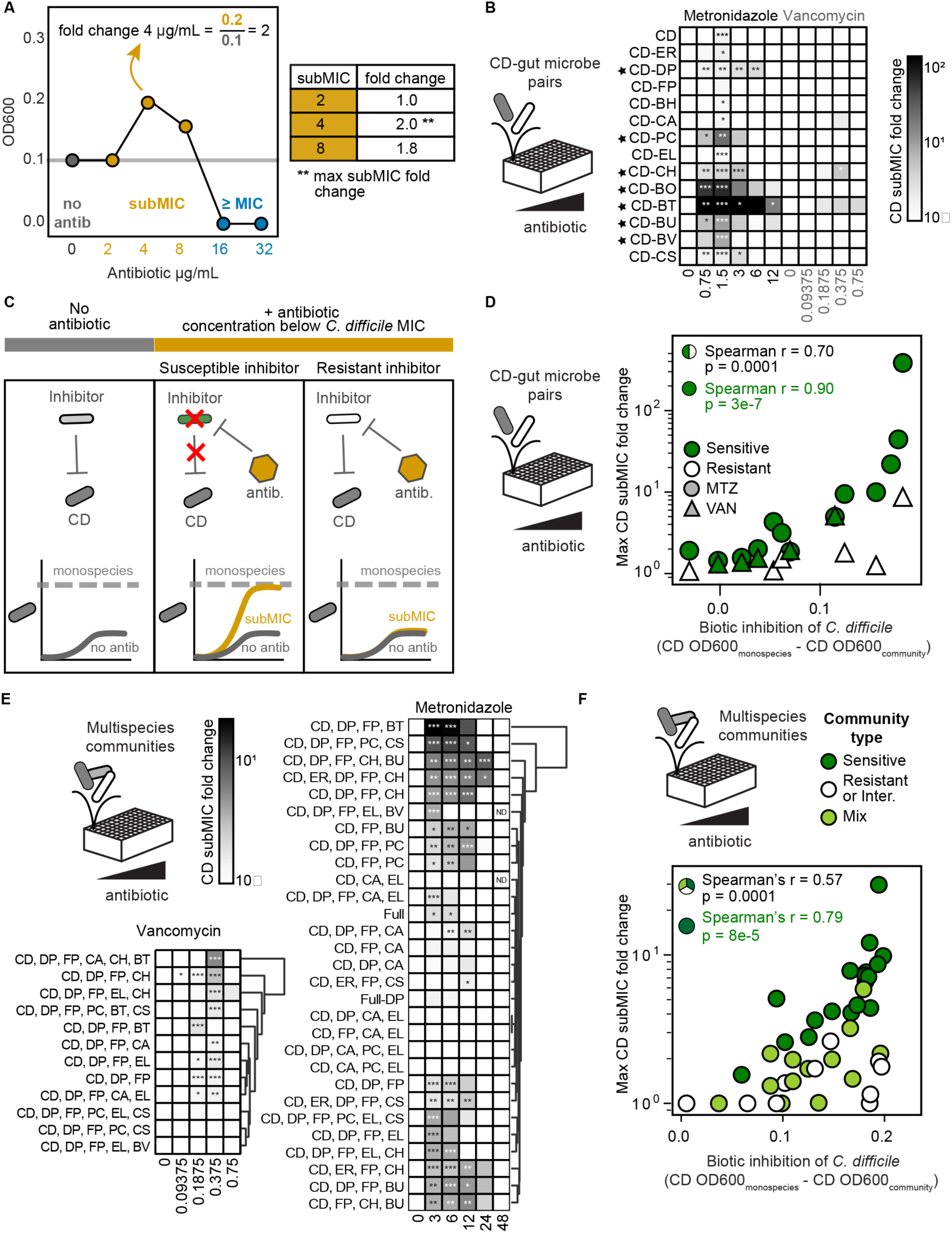
*C. difficile* growth is enhanced in communities with antibiotic-sensitive biotic inhibitors in response to low antibiotic concentrations. **(a)** Schematic of subMIC fold change metric. The x-axis is semi-log scale. Table indicates subMIC fold changes for the three subMICs (2, 4, and 8 µg/mL) in this example. Maximum subMIC fold change (indicated by **) is defined as the largest fold change across all subMICs. **(b)** Heatmap of subMIC fold changes for *C. difficile* in pairwise communities at each concentration at 48 hour timepoint. Shading indicates subMIC fold change. SubMIC fold change is calculated as the average *C. difficile* absolute abundance at the subMIC divided by the average *C. difficile* absolute abundance in the absence of antibiotic, where the average is of n=1 to n=8 biological replicates. Asterisks indicate significant difference (*P < 0.05, **P < 0.01, ***P < 0.001) between *C. difficile* OD600 at subMIC and *C. difficile* OD600 in the absence of antibiotic according to an unpaired t-test. Black stars indicate pairs with a maximum subMIC fold change significantly greater than the maximum subMIC fold change of *C. difficile* monospecies (p<0.05 by an unpaired t-test). **(c)** Schematic of *C. difficile* growth in the presence of antibiotic sensitive and resistant inhibitors. **(d)** Scatterplot of maximum *C. difficile* subMIC fold change in pairs as a function of biotic inhibition of *C. difficile* in pairs for metronidazole (MTZ, circles) and vancomycin (VAN, triangles). Biotic inhibition of *C. difficile* (x-axis) is equal to the average *C. difficile* OD600 in monospecies minus the average *C. difficile* OD600 in the pairwise community in no antibiotic conditions. The maximum *C. difficile* subMIC fold change (y-axis) is the maximum of the *C. difficile* subMIC fold change at all subMICs at 48 hours. The subMIC fold change is calculated as in panel B, with n=1 to n=8 biological replicates. Species are classified as sensitive if the species monospecies MIC was less than the monospecies MIC of *C. difficile* and resistant if the species monospecies MIC was greater than or equal to the monospecies MIC of *C. difficile*. Each point represents one pair in metronidazole or vancomycin. Spearman correlation is annotated for all data points (black) and for pairwise communities with sensitive gut microbes only (green). **(e)** Heatmap of subMIC fold changes for *C. difficile* in multispecies communities at each concentration of metronidazole and vancomycin at 48 hours. Shading indicates subMIC fold change. SubMIC fold change is calculated as in panel B, with n=1 to n=4 biological replicates. Asterisks indicate significant difference (*P < 0.05, **P < 0.01, ***P < 0.001) between *C. difficile* OD600 at subMIC and *C. difficile* OD600 in the absence of antibiotic according to an unpaired t-test. Communities clustered by subMIC fold change using Euclidean distance hierarchical clustering. “Full” indicates 14-member community, “Full-DP” indicates 13-member community containing all species except DP. **(f)** Scatterplot of maximum *C. difficile* subMIC fold change in multispecies communities as a function of biotic inhibition of *C. difficile* in multispecies communities for metronidazole (circles) and vancomycin (triangles). Biotic inhibition of *C. difficile* (x-axis) is calculated as in panel D. The maximum *C. difficile* subMIC fold change (y-axis) is calculated as in panel B, with n=1 to n=4 biological replicates. Community type is sensitive (“Sens.”) if all species excluding *C. difficile* are sensitive inhibitors. Community type is resistant or intermediate (“Resis. or Intermed.”) if all species excluding *C. difficile* are either intermediate-inhibitors or resistant-inhibitors. Community type is considered “mix” if the given community contains both sensitive inhibitors and resistant- or intermediate-inhibitors. Species are categorized as biotic inhibitors if the absolute abundance of *C. difficile* in the presence of this species was significantly lower than the absolute abundance of *C. difficile* in monospecies, in the absence of antibiotics, as determined by an unpaired t-test. Species are classified as sensitive and resistant as in panel D. *C. aerofaciens* was classified as intermediate (see text). Each point represents one community in the presence of metronidazole or vancomycin. Spearman correlation annotated for all data points (black) and for sensitive communities only (green).

In monoculture, *C. difficile*’s growth was moderately enhanced in the presence of a single subMIC (1.5 µg/mL metronidazole, subMIC fold change of 1.6, **Supplementary Figure 2A**). Time-series OD600 measurements of *C. difficile* monoculture revealed that this moderate growth enhancement was due to a difference in growth phase in the presence and absence of the antibiotic at the measured timepoint (48 hours) (**Supplementary Figure 2CD**). By contrast, *C. difficile*’s growth was enhanced in two pairwise communities in the presence of subMICs compared to no treatment over multiple time points that was not due to differences in growth phase (**Supplementary Figure 2EF**). Thus, while differences in monoculture growth phases yielded a modest enhancement of *C. difficile* in response to a single subMIC, a different mechanism determined the observed larger and sustained growth enhancements in pairwise communities.

We hypothesized the action of the antibiotic in certain pairwise communities could relieve the biotic inhibition of *C. difficile*, which in turn enhances *C. difficile* growth. This would occur in pairwise communities with competitors (i.e., biotic inhibitors) with a higher susceptibility to the antibiotic than *C. difficile*. By contrast, if a biotic inhibitor displayed a higher resistance to the antibiotic than *C. difficile,* the degree of biotic inhibition would remain unchanged at subMICs and thus *C. difficile*’s growth would not change substantially compared to its growth in absence of the antibiotic (**Figure 3C**). Supporting this hypothesis, the maximum subMIC fold change of *C. difficile* displayed a strong positive correlation with the degree of biotic inhibition in the presence of the co-culture partner species (**Figure 3D**). In addition, the correlation was enhanced in pairwise communities containing gut microbes that were sensitive to the antibiotic (**Figure 3D**). In the presence of both individual antibiotics, the maximum subMIC fold change of *C. difficile* was significantly higher when paired with biotic inhibitors with higher sensitivity to the antibiotic than *C. difficile* (**Supplementary Figure 2G**). In sum, antibiotic induced inhibition of highly susceptible biotic inhibitors enhanced *C. difficile*’s growth in response to subMICs. This suggests that subMICs, which can occur during clinical dosing regimens (at the beginning and end of regimens and between daily dosages^25^), could enhance *C. difficile*’s growth if patients harbor antibiotic-sensitive biotic inhibitors in their gut microbiota.

### Enhancement of C. difficile abundance in multi-species communities in the presence of low antibiotic concentrations

We investigated if the trends observed in pairwise communities persist in multi-species communities that are more representative of the human gut microbiota. The communities consisted of 2- and 3-member core communities (CC) predicted to display minimal biotic inhibition of *C. difficile* guided by a previously developed dynamic computational model of our system^4^ (CC-2: *D. piger* and *Faecalibacterium prausnitzii*. CC-3: *D. piger*, *F. prausnitzii*, and *Eggerthella lenta*). In addition to this core community, we introduced at least one antibiotic-sensitive inhibitor, antibiotic-resistant inhibitor, or a combination of species in these two groups, creating 10 communities for metronidazole and 12 communities for vancomycin (3- to 6-members). To further explore the behavior of communities in response to antibiotic perturbations, we characterized the response of 3-, 4-, 5-, 13-, or 14-member communities (19 total) with no consistent core members to metronidazole.

In 5 of 29 communities examined in the presence of metronidazole, *C. difficile*’s MIC was 4-fold greater than in monoculture, in one community the MIC was 4-fold lower, and the remaining communities displayed moderate (2-fold, 13 communities) or no change (10 communities) (**Supplementary Figure 3A, Supplementary Table 1**). In 4 of 12 communities examined in the presence of vancomycin, *C. difficile’s* growth was strongly inhibited and no MIC could be calculated. Of the remaining 8 communities, 7 displayed no change in *C. difficile*’s MIC and *C. difficile* displayed a moderate (2-fold) increase in MIC in one community (**Supplementary Figure 3B, Supplementary Table 2**). Consistent with the pairwise community data, *C. difficile’s* metronidazole MIC was altered in certain communities, but *C. difficile’s* vancomycin MIC was not substantially altered in any of the characterized communities.

We tested if the multi-species communities containing antibiotic-sensitive biotic inhibitors displayed an enhancement in *C. difficile*’s growth at subMICs. In response to metronidazole or vancomycin, *C. difficile’s* growth was enhanced in response to at least one subMIC in most communities (21 of 29 communities for metronidazole, 9 of 12 communities for vancomycin, **Figure 3E**, **Supplementary Figure 4AB**). Consistent with the pairwise community data, the degree of biotic inhibition of *C. difficile* was positively correlated with the magnitude of the maximum subMIC fold change (**Figure 3F**). In addition, the correlation was stronger for communities composed of only antibiotic-sensitive biotic inhibitors (**Figure 3F**). In addition, the number of sensitive inhibitors in the community and the absolute abundance of sensitive inhibitors displayed a positive correlation with the maximum subMIC fold change (**Supplementary Figure 5A**). Communities containing antibiotic-sensitive biotic inhibitors had a significantly higher average subMIC fold change than communities containing only resistant inhibitors or a mix of both inhibitor types (**Supplementary Figure 5B**).

Communities containing a mix of antibiotic sensitive and resistant inhibitors, mirroring the makeup of the gut microbiota, displayed low subMIC fold changes similar in magnitude to resistant biotic inhibitor communities (**Supplementary Figure 5B**). In communities with a non-zero abundance of resistant biotic inhibitors, the correlation between the growth enhancement of *C. difficile* and the number of sensitive inhibitors or abundance of sensitive inhibitors vanished (**Supplementary Figure 5A**). These data suggest that resistant biotic inhibitors can suppress the

*C. difficile* growth enhancement caused by sensitive biotic inhibitors. We analyzed if the growth enhancement of *C. difficile* in multi-species communities could be predicted based on the sum of the growth enhancements of *C. difficile* in pairwise communities. While the relationship displayed a moderate correlation, it was not additive (**Supplementary Figure 5CD**). This demonstrates that community effects, such as inter-species interactions between constituent community members and growth enhancement suppression by resistant inhibitors, play key roles in determining the magnitude of growth enhancement of *C. difficile* at subMICs.

Since *C. difficile* growth enhancement at subMICs can also occur in multi-species communities, this response could also occur in the human gut microbiome. Our results suggest that either narrow spectrum antibiotics which inhibit the fewest members of the microbiota or microbiome interventions that enhance the abundance of antibiotic resistant inhibitors would limit the growth enhancement of *C. difficile* in response to subMICs.

### A dynamic ecological model representing pairwise interactions and monospecies antibiotic susceptibilities can capture the response of C. difficile to subMICs

We hypothesized that a model capturing the dynamics of species growth, inter-species interactions and monospecies antibiotic susceptibilities could recapitulate the observed trends in *C. difficile* growth enhancement at subMICs. We assumed that inferred inter-species interactions in the absence of antibiotics contributed to the community response to antibiotics. We tested whether a model that neglects complex antibiotic-dependent inter-species interactions could predict the qualitative trends of *C. difficile* growth at subMICs.

The gLV model is a system of coupled ordinary differential equations which captures individual species’ growth rate and pairwise interactions with each community member. We use an expansion of the generalized Lotka-Volterra model (gLV) that captures antibiotic perturbations^28^ (**Figure 4A**). In this expanded model, the growth of each species is modified by an antibiotic term consisting of the concentration of antibiotic and the susceptibility of each species (B_i_) (**Methods**, **Figure 4A**).

**Figure 4:**
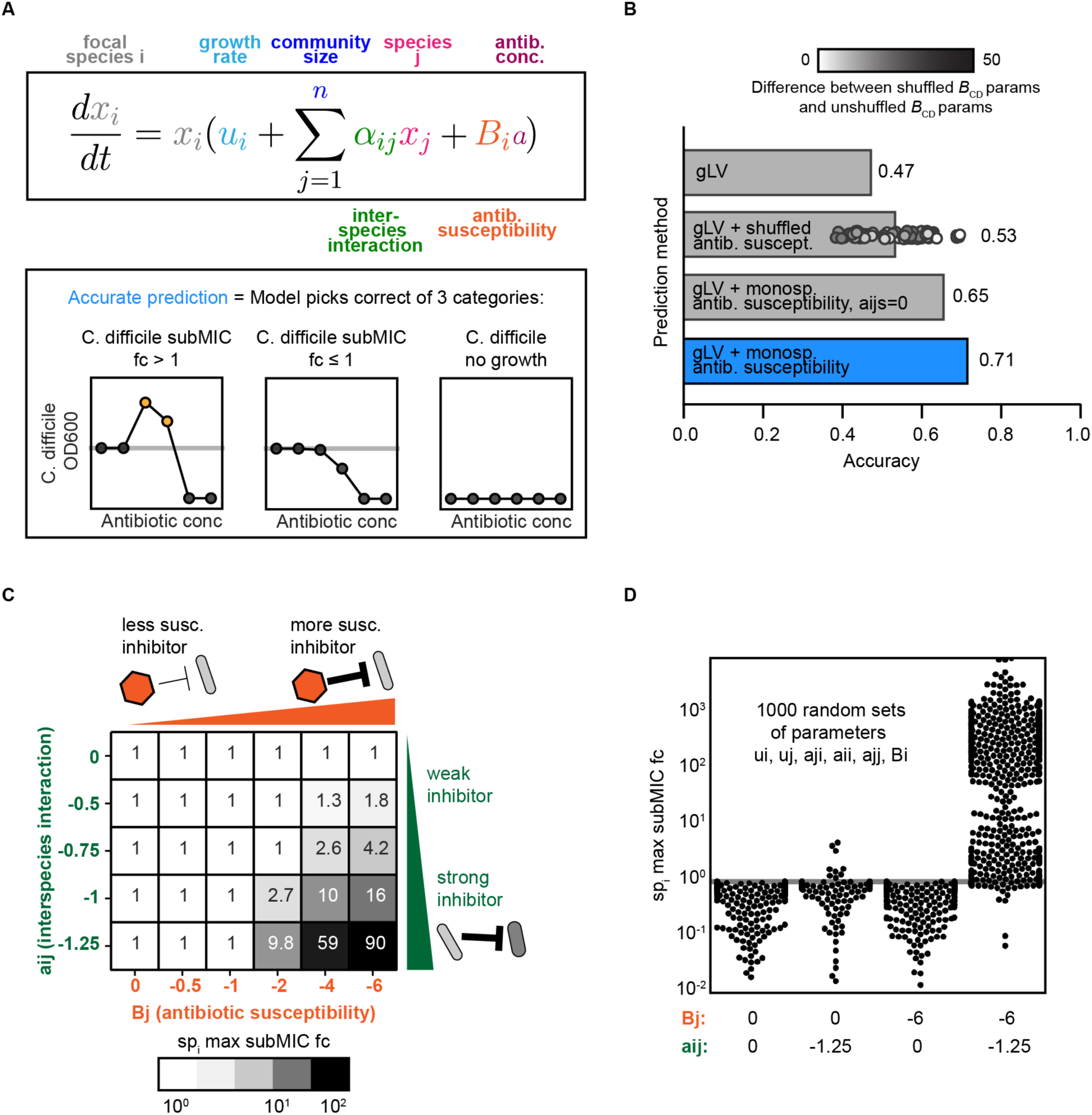
A modified generalized Lotka-Volterra model of community response to antibiotics captures antibiotic sensitive biotic inhibitor trend. **(a)** Top: Schematic of the antibiotic expansion of the gLV model. Bottom: Schematic of accuracy metric used in panel B. **(b)** Qualitative accuracy of multiple models for pairwise and multispecies communities. Models include 1) Standard gLV model lacking an antibiotic term; 2) gLV model with randomly shuffled antibiotic susceptibility parameters (average of 100 permutations). 100 shuffled values shown as points, shaded according to the absolute value of the difference between the shuffled *C. difficile* antibiotic susceptibility parameter and the original *C. difficile* antibiotic susceptibility parameter; 3) gLV model with antibiotic susceptibility terms inferred from monospecies with all interaction parameters set to zero (a_ij_ = 0, where *i* != *j*); 4) full gLV model with antibiotic susceptibility terms inferred from monospecies. **(c)** Simulated maximum subMIC fold changes for a focal species in a pairwise community for 48 hours for a representative parameter set. Growth rates of the two species are equal (u_i_ = u_j_ = 0.25), intra-species interactions are equal (a_ii_ = a_jj_ = -0.8), and initial ODs are equal (x_i_(0) = x_j_(0) = 0.0022). A_ji_ is set to 0 and B_i_ is set to -2. **(d)** Simulated max subMIC fold change for a focal species in a pairwise community for 1000 randomly sampled parameter sets. Parameters *B*_j_ and a_ij_ are set to constant values of 0 or -6 and 0 or -1.25 respectively. Other parameters were randomly sampled between upper and lower bounds. The bounds on each parameter value include a_ji_ (-1.25, 1.25), growth rates (0,1), intraspecies interactions (-1.25, 0), *B*_i_ (-6,0). Gray horizontal line at y=1 indicates no change in growth compared to the no antibiotic condition.

The monospecies antibiotic susceptibility parameters were inferred from measurements of individual species growth in the presence of a range of antibiotic concentrations (2-fold dilutions) (Methods, **Supplementary Figure 1**). We used growth rate and inter-species interaction parameters that were previously inferred based on absolute abundance measurements of 159 monospecies and communities (2 to 14-member) in the absence of antibiotics^4^.

We evaluated whether this model could qualitatively predict the trends in growth enhancement of *C. difficile* in the subMIC range (**Supplementary Figure 6, 7**). *C. difficile*’s response to each antibiotic concentration was classified into three categories (*C. difficile* subMIC fold-change > 1, *C. difficile* subMIC fold-change ≤ 1, or no growth of *C. difficile,* **Figure 4A**). The expanded gLV model correctly predicts *C. difficile*’s qualitative response to 71% of antibiotic concentrations (**Figure 4B**, **Supplementary Figure 8A**). We designed a set of null models to evaluate the contribution of different terms of the expanded gLV model to model performance. The full model displays higher accuracy than a null model that lacked antibiotic susceptibility terms or had randomly shuffled antibiotic susceptibility parameters (47% and 53% accuracy, **Figure 4C**). These data highlight that the fitted monospecies susceptibilities play a major role in the model’s predictive performance. The full model also outperformed a null model that lacked inter-species interaction terms (65%, **Figure 4C**), indicating that the inferred inter-species interactions contribute to the growth enhancement of *C. difficile* in response to subMICs. However, the monospecies antibiotic susceptibilities are a dominant driver of this behavior. Taken together, these findings indicate that biotic inhibition and monospecies antibiotic susceptibility are major variables determining the growth of *C. difficile* in response to subMICs in microbial communities. We explored model simulations to determine if the enhancement of *C. difficile* growth at subMICs required a sensitive biotic inhibitor in a wide variety of communities, beyond those that were experimentally characterized. To this end, we simulated 1000 pairwise communities with a wide range of growth rates, inter-species interactions, and antibiotic susceptibilities. When paired with a sensitive biotic inhibitor, growth is enhanced at subMICs, consistent with the trends observed in our experiments (**Figure 4D**). By contrast, growth enhancements are not observed in simulated communities that lack an antibiotic-sensitive biotic inhibitor of the focal species (**Figure 4D**). This trend is also present in larger communities (**Supplementary Figure 8B**). The focal species’ growth enhancement at subMICs increases with biotic inhibition and antibiotic susceptibility of the inhibitor species (**Figure 4E**). Overall, these model simulations suggest that the antibiotic sensitive inhibitor trend we observe with *C. difficile* in human gut communities is generalizable to other species and communities with variable richness, interaction networks, and antibiotic susceptibilities.

*D. piger substantially increases C. difficile’s minimum inhibitory concentration to metronidazole* Changes in antibiotic tolerance of a pathogen due to significant microbial interactions could reduce the efficacy of antibiotic treatments. In the presence of *D. piger*, we observed a substantial increase in *C. difficile* MIC, in addition to a moderate growth enhancement of *C. difficile* at subMICs (**Figure 2B**). Consistent with the proposed antibiotic-sensitive biotic inhibition mechanism, *D. piger* was a weak biotic inhibitor of *C. difficile* and was more sensitive to metronidazole than *C. difficile* (**Figure 2A, 3C**). However, the substantial increase in *C. difficile* MIC in the presence versus absence of *D. piger* (≥24 µg/mL compared to 6 µg/mL) was not explained by the proposed antibiotic-sensitive biotic inhibition mechanism and the expanded gLV antibiotic model failed to predict this trend (**Supplementary Figure 6A**). While competitive interactions (i.e. resource competition or biological warfare) contributed to the observed subMIC growth response, the mechanism of increased tolerance of *C. difficile* to metronidazole in the presence of *D. piger* was unknown.

We investigated the robustness of this trend across different environmental conditions and for different *C. difficile* isolates. *C. difficile* displayed increased tolerance in co-culture with *D. piger* and in monoculture in *D. piger*’s spent media (≥8-fold increase in MIC) (**Figure 5A**). These data indicate that the increase in *C. difficile* metronidazole tolerance was not dependent on cell-to-cell contact with *D. piger* or prior exposure of *D. piger* to metronidazole. *C. difficile* displayed higher metronidazole tolerance in *D. piger* spent media harvested at late exponential phase/early stationary phase than in spent media harvested at earlier time points in exponential phase (**Supplementary Figure 9AB**). Further, multiple *C. difficile* clinical isolates displayed enhanced tolerance in *D. piger* spent media (**Supplementary Figure 9C**). Finally, *C. difficile* exhibited enhanced metronidazole tolerance in a different chemically defined growth medium (**Supplementary Figure 9D**). These data demonstrate that the protective effect of *D. piger* on *C. difficile’s* response to metronidazole was robust to variations in *C. difficile*’s strain background, media composition and the growth phase of *D. piger*.

**Figure 5:**
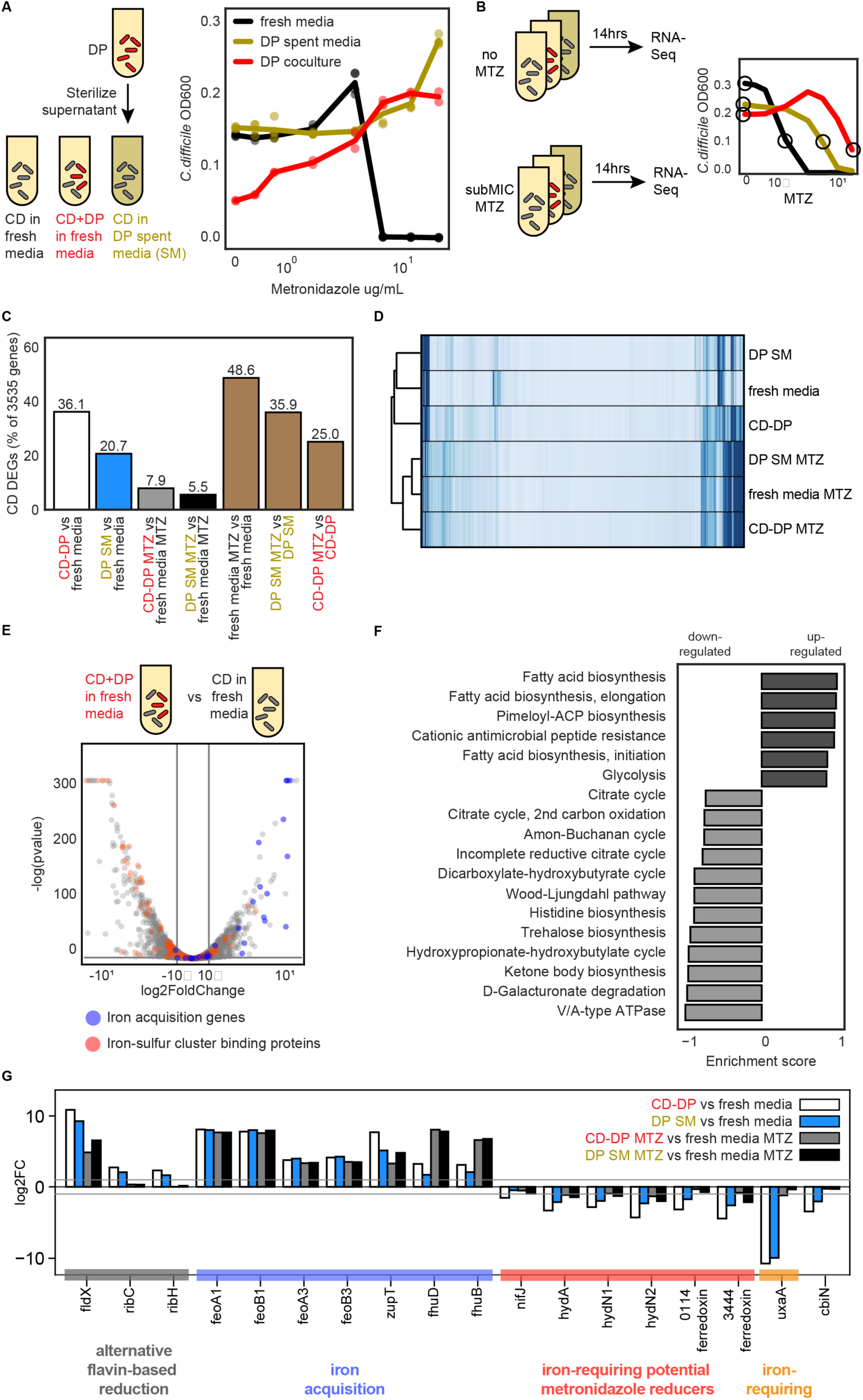
*D. piger* increases *C. difficile* tolerance to metronidazole and induces metal starvation genome-wide transcriptional response. **(a)** Abundance of *C. difficile* at 41 hours in response to different metronidazole concentrations. The OD600 value for *C. difficile* in the presence of *D. piger* is calculated OD600 (OD600 multiplied by relative abundance from 16S rRNA gene sequencing). The x-axis is semi-log scale. Data points represent biological replicates. Lines indicate average of n=2 to n=4 biological replicates. **(b)** Schematic of genome-wide transcriptional profiling experiment. The x-axis is semi-log scale of metronidazole (MTZ) concentration. Lines indicate average *C. difficile* OD600 at 14 hours for n=4 biological replicates. Circles indicate conditions sampled for RNA-Seq. (c) *C. difficile* differentially expressed genes (DEGs) between culture conditions. DEGs are defined as genes that displayed greater than 2-fold change and a p-value less than 0.05. **(d)** Clustered heatmap of reads per kilobase million (RPKM) for each gene (rows) and in each sample (columns) for *C. difficile*. Each column represents the average of n=2 biological replicates. Hierarchical clustering was performed based on Euclidean distance using the single linkage method of the python SciPy clustering package. **(e)** Volcano plot of log transformed transcriptional fold changes for *C. difficile* in the presence of *D. piger.* Gray vertical lines indicate 2-fold change and the gray horizontal line indicates the statistical significance threshold (p=0.05). Blue indicates genes annotated to be involved in iron or zinc import. Red indicates genes predicted to contain iron-sulfur clusters by MetalPredator^29^. **(f)** Enriched gene sets in *C. difficile* grown in the presence of *D. piger* compared to *C. difficile* grown in fresh media. All gene sets with significant enrichment scores from Gene Set Enrichment Analysis (GSEA) are shown. Gene sets are defined using modules from the Kyoto Encyclopedia of Genes and Genomes (KEGG). **(g)** Bar plot of the log transformed fold changes of a set of genes across different conditions. Gray horizontal lines indicate 2-fold change.

*D. piger* could alter *C. difficile’*s metronidazole tolerance via a direct chemical interaction with metronidazole which reduces its efficacy (*e.g.,* degradation or sequestration) or an indirect effect that modifies *C. difficile*’s intracellular networks which in turn yields an increase in metronidazole tolerance. To test whether there was a direct chemical interaction, we incubated metronidazole in either *D. piger* spent media or fresh media and characterized *C. difficile*’s growth response to each of these conditions in fresh media. *C. difficile*’s tolerance to metronidazole was similar in these conditions, indicating that compounds present in *D. piger*’s spent media do not directly interact with metronidazole to reduce its activity (**Supplementary Figure 9E**).

### C. difficile displays a genome-wide transcriptional signature of iron-limitation in response to D. piger

We considered indirect effects of *D. piger* on *C. difficile*’s intracellular network activities which in turn could alter *C. difficile*’s response to metronidazole. We first considered resource competition as a possible indirect effect because resource competition is a common mechanism driving inter-species interactions in microbial communities. *C. difficile*’s abundance was not substantially lower in *D. piger*’s spent media than monoculture (**Figure 5A**), suggesting that resource competition was not a major driver of the increased tolerance. In addition, supplementing concentrated fresh media into the *D. piger* spent media, which would alleviate any resource competition, did not alter *C. difficile’s* metronidazole tolerance (**Supplementary Figure 9F**). Therefore, these data suggest that *D. piger* did not affect metronidazole tolerance via resource competition.

We considered if *D. piger* affected the metronidazole tolerance of *C. difficile* by altering patterns in gene expression. To this end, we performed genome-wide transcriptomic profiling of *C. difficile* in late exponential phase in the following environments: (1) monoculture in fresh media, (2) monoculture in *D. piger* spent media, and (3) co-culture with *D. piger*. Each of these cultures was exposed to a subMIC of metronidazole that reduced growth by approximately 50% or no treatment was applied (**Figure 5B**). The antibiotic concentration was chosen to allow sufficient growth for transcriptional profiling.

*C. difficile*’s gene expression profile in *D. piger* conditions (i.e., *D. piger* spent media and co-culture with *D. piger*) was significantly altered compared to *C. difficile* monoculture (**Table 1**, **Supplementary Table 3**). Differential expressed genes (DEGs) were defined as genes with >2-fold change and a p-value less than 0.05. In the absence of antibiotics, 36% and 21% of *C. difficile*’s 3535 genes were differentially expressed in the *D. piger* co-culture and spent media, respectively, compared to the *C. difficile* monoculture in fresh media (**Figure 5C**, white and blue bars). Of these genes, 72 in the *D. piger* co-culture and 47 in the *D. piger* spent media exhibited >8-fold change, demonstrating that the presence of *D. piger* had a large impact on *C. difficile*’s gene expression profile. In the presence of metronidazole, a smaller percentage of genes were differentially expressed between the *D. piger* and fresh media conditions (8% and 6%, **Figure 5C**, gray and black bars). Large shifts in *C. difficile* gene expression occurred due to the addition of metronidazole, with 25-49% of genes differentially expressed (**Figure 5C**, brown bars). The three metronidazole conditions clustered closely together, indicating that the antibiotic induced similar changes in gene expression regardless of the media condition or ecological context (**Figure 5D**).

**Table 1:**
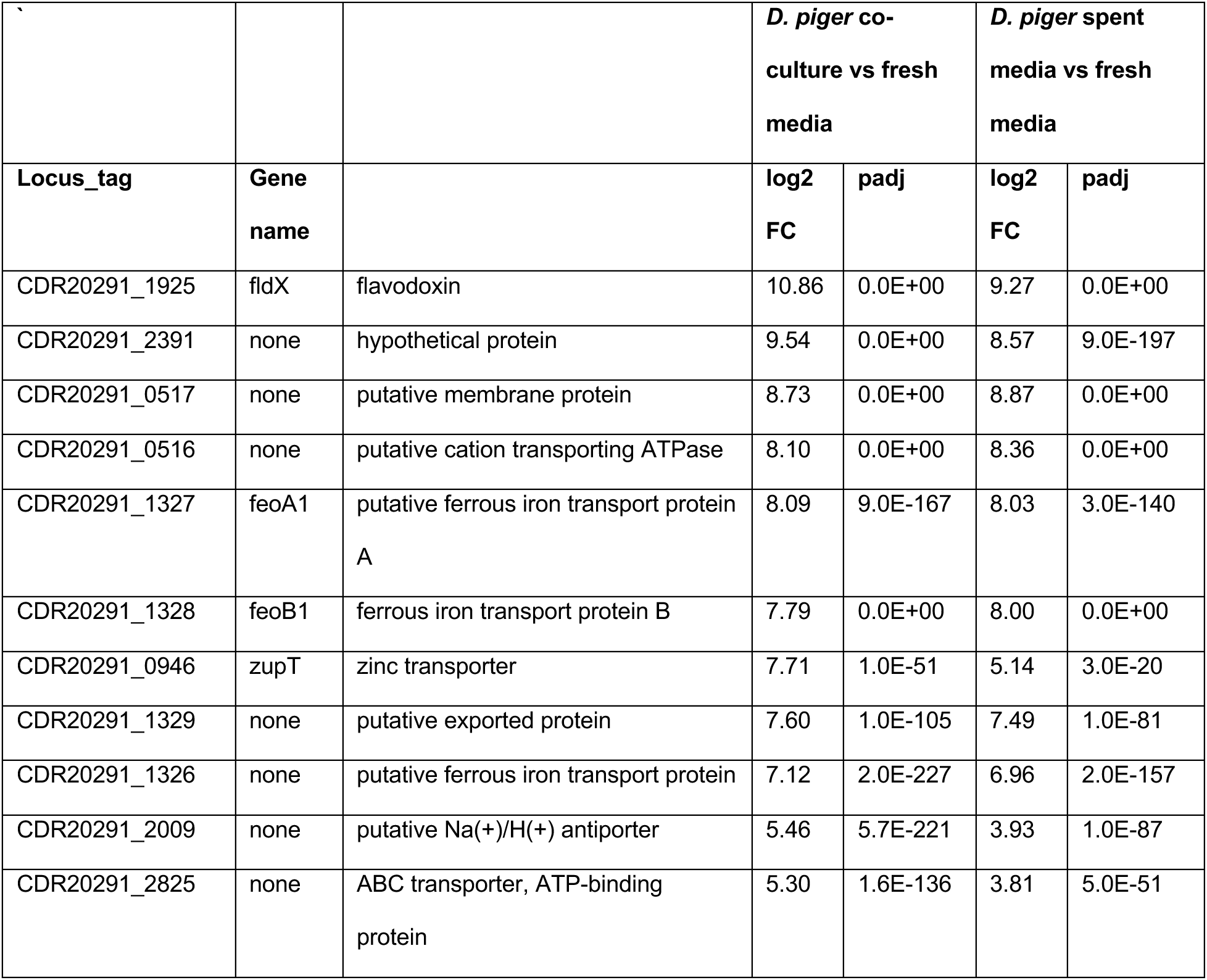
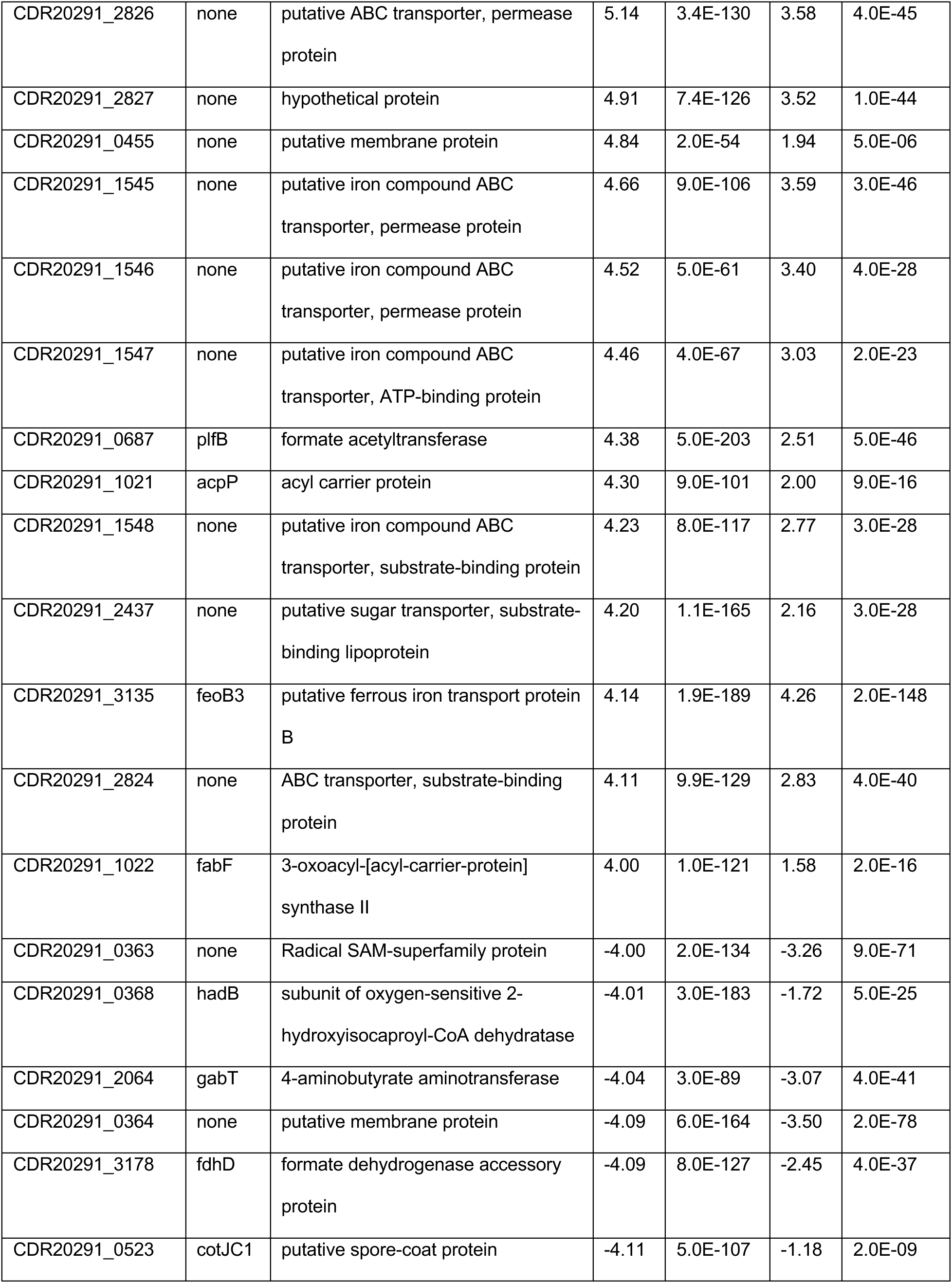
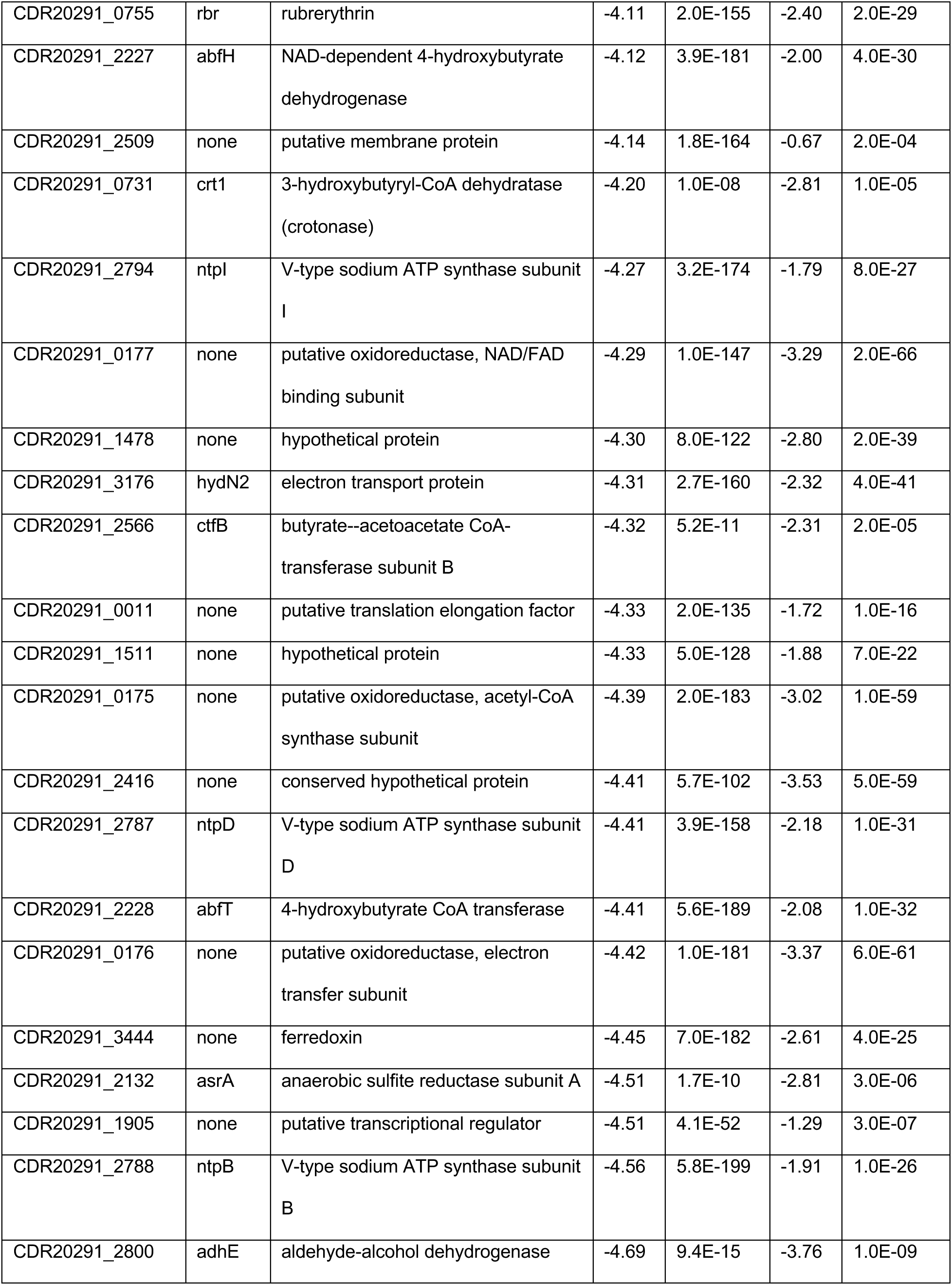
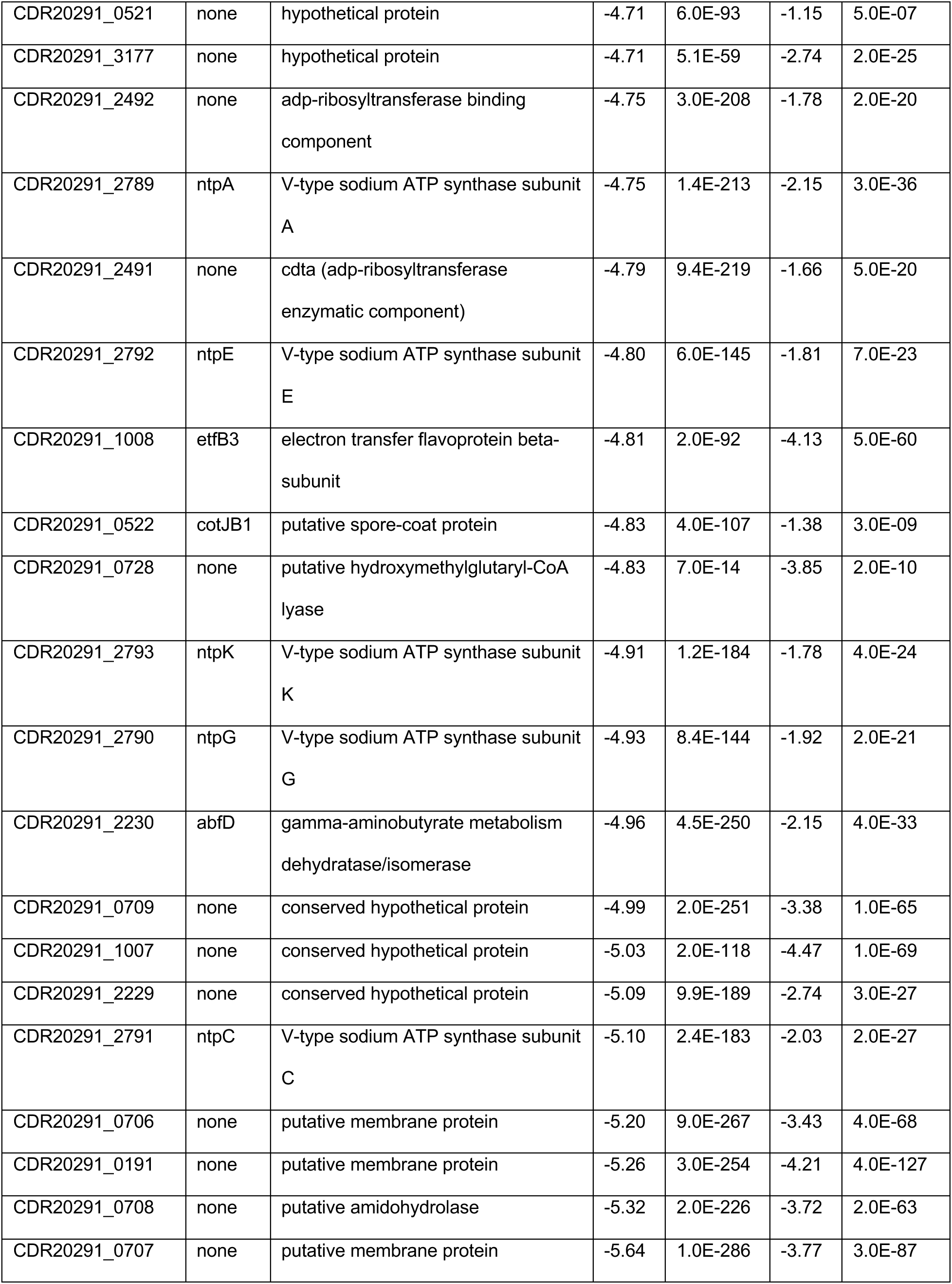
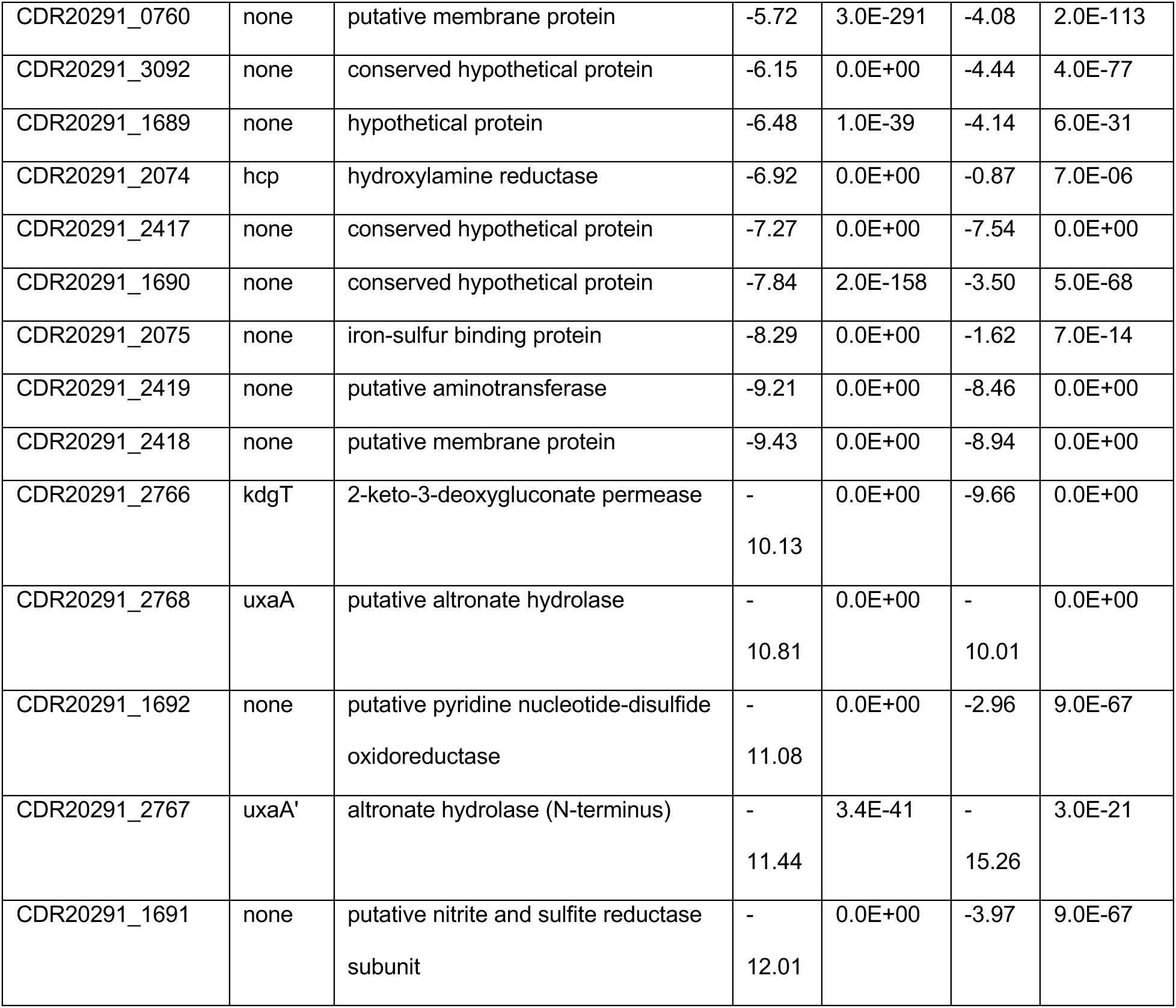
*C. difficile* gene expression in *D. piger* conditions compared to fresh media for genes with >16-fold differential expression.

To provide deeper insights into specific genes that varied across conditions, we analyzed the genes with the largest expression changes in *D. piger* conditions since these conditions displayed substantial shifts in gene expression. Many of the genes upregulated in *D. piger* and *D. piger* antibiotic conditions compared to fresh media had annotated roles in iron acquisition (blue points in **Figure 5E**, **Supplementary Figure 10**). These data suggest that *D. piger* created an iron-limited environment for *C. difficile.* To acquire iron, *C. difficile* uses iron permeases to import free ferrous iron into the cell^30, 31^. We observed >10-fold upregulation in two ferrous transporters (*feoAB1, feoAB3*) in *D. piger* and *D. piger* metronidazole conditions (**Table 1**, **Figure 5G**). Alternative transporters were also upregulated, such as uptake genes for ferrichrome siderophores (*fhuB, fhuD)* and catecholate siderophores (*CDR20291_1545 – 1548*, orthologs of *yclNOPQ* in CD630) as well as a zinc-transporter gene (*zupT*). ZupT may be a transporter for iron as well as zinc, which has been observed previously in other bacterial species^32^. These iron acquisition genes (*feo, fhu, ycl,* and *zupT*) are regulated by the iron regulator *fur*^30^ and have been shown to be upregulated in response to iron starvation^33, 34^, suggesting that iron limitation in *D. piger* conditions was responsible for the observed changes. Consistent with the iron limitation signature, ferrous transporters that were not up-regulated (*feoAB2*) have been documented as unresponsive to iron-limitation^30, 33^.

In addition to genes directly related to iron transport, many of the differentially expressed genes in *C. difficile* were linked to iron-metabolism. Flavodoxin (*fldX*) was the most upregulated gene in the *D. piger* conditions (**Table 1, Figure 5G**). Flavodoxin can replace iron-requiring electron transfer proteins (such as ferredoxin) in iron-limiting conditions^35^. Flavodoxin uses a different uses a non-iron cofactor, flavin, whose biosynthesis genes (*ribACH*) were upregulated in *D. piger* conditions (**Table 1**). Previous studies have shown that flavodoxin was regulated by *fur* and de-repressed in iron-limiting conditions^30, 33, 34^ and riboflavin biosynthesis genes were upregulated in iron-limiting conditions^34^. Iron-containing proteins are also connected to iron-metabolism and were down-regulated in iron-limited conditions^34^. Using a bioinformatic analysis, 9% of *C. difficile’s* downregulated genes (62 of 678 genes) in the *D. piger* co-culture condition are predicted to contain iron-sulfur clusters (red points in **Figure 5E**, **Supplementary Table 4**). In sum, genes for replacing iron-requiring ferredoxin were up-regulated, whereas transcripts for iron-requiring proteins were down-regulated, consistent with an iron-limited environment for *C. difficile* created by *D. piger*.

To identify other biological pathways that were significantly altered in *D. piger* conditions, we performed gene set enrichment analysis using KEGG modules (**Figure 5F**, **Supplementary Figure 10**). Many of the identified pathways could be connected to iron. For example, the majority of the enzymes in the downregulated Wood-Ljungdahl pathway have iron-sulfur clusters, namely carbon monoxide dehydrogenase (*cooCS, CDR20291_0653, CDR20291_0655)*, [NiFe] hydrogenase (*hydAN1N2*), and formate dehydrogenase H (*fdhF*)^36–38^. These enzymes have been shown to be down-regulated under iron-limited conditions^34^. Pathways for cationic antimicrobial peptide resistance and fatty-acid biosynthesis were upregulated, which has been previously attributed to iron-limitation^34^. Similarly, the downregulation of pathways for V-type ATPases, D-galacturonate degradation, and trehalose biosynthesis have been connected to iron-limitation (**Figure 5F**)^34^. *D. piger* also affected the expression of *C. difficile* toxins (*tcdA, tcdB*, and binary toxin genes *CDR20291_2491* and *CDR20291_2492*) which were downregulated between 4- and 27-fold in the *D. piger* conditions (**Supplementary Table 3**).

To provide further insights into the contribution of iron limitation on the patterns of gene expression in *C. difficile*, we evaluated the relationship between gene expression changes in *C. difficile* in the presence of *D. piger* and gene expression changes in *C. difficile* in iron-limited media previously characterized in a separate study^34^. For all differentially expressed genes, we compared the log2 fold changes between *C. difficile* in the *D. piger* co-culture and *C. difficile* in monoculture in the absence of metronidazole with the log2 fold changes observed between *C. difficile* in iron-limited and iron-rich media in the previous study. These fold changes displayed an informative relationship and were qualitatively consistent for 89% of genes between the two studies (**Supplementary Figure 11A**, Pearson r= 0.61, p=8*10^-54^). Similarly, the *D. piger* spent media showed 87% qualitative agreement (**Supplementary Figure 11B**). The informative relationship between these datasets suggests that the shifts in gene expression for the majority of *C. difficile*’s genes in the *D. piger* conditions can be explained by iron limitation.

Overall, these data suggest that *D. piger* created an iron limited environment for *C. difficile*, as iron acquisition genes and alternative genes involved in flavin metabolism were upregulated whereas transcripts for iron-requiring proteins were downregulated (**Figure 5G**). These trends were observed in both *D. piger* co-culture and *D. piger* spent media and were also consistent with the trends in the *D. piger* conditions with metronidazole, although there were a smaller number of differentially expressed genes in the presence of the antibiotic (**Figure 5G**).

### Differentially expressed genes in C. difficile in the presence of D. piger are linked to metronidazole resistance

Since the majority of *C. difficile’s* differentially expressed genes could be explained by iron limitation, we investigated the connection between iron limitation and *C. difficile* metronidazole tolerance. To identify potential connections, we compared the set of differentially expressed genes in the presence of *D. piger* to genes previously shown to play a role in metronidazole resistance in *C. difficile*.

In two studies of metronidazole resistant *C. difficile* mutants, iron-related genes were implicated in metronidazole resistance. In a study of *C. difficile* 630, multiple mutants all acquired a truncation in *feoB1,* which resulted in reduced intracellular iron and a shift to flavodoxin^39^. Similarly, in a R20291 mutant, iron uptake genes were downregulated and a shift to flavodoxin was observed^40^. In each of these studies, the proposed mechanism of metronidazole resistance was attributed to down-regulation of enzymes predicted to reduce metronidazole to its active form. In the *D. piger* conditions, enzymes hypothesized to reduce metronidazole were also down-regulated, namely ferredoxin genes (*fdxA, CDR20291_0114, CDR20291_3444*), pyruvate-ferredoxin/flavodoxin oxidoreductase (PFOR) (*nifJ*), and hydrogenases (*hydA*, *hydN1*, *hydN2*) (**Table 1, Figure 5G**)^10, 11^. Down-regulation of these enzymes in *D. piger* conditions may reduce the rate of conversion of metronidazole into its active form, thus enhancing the tolerance of *C. difficile*. The downregulated enzymes are predicted to contain iron clusters (**Supplementary Table 4**) suggesting that their downregulation and the subsequent increase in metronidazole tolerance could be attributed to iron limitation.

This mechanism of resistance has been proposed in other species beyond *C. difficile*, such as *Bacteroides fragilis* metronidazole resistant mutants that displayed reduced PFOR expression^11^. Therefore, if this mechanism was responsible for the enhancement in *C. difficile*’s metronidazole tolerance by *D. piger*, *Bacteroides* species should also display enhanced tolerance. Indeed, *B. thetaiotaomicron* displayed an increased tolerance to metronidazole in *D. piger* spent media compared to fresh media (**Supplementary Figure 12**).

We also identified that *cbiN,* a putative cobalt transporter that was downregulated in the *D. piger* conditions (**Table 1, Figure 5G**) and in iron-limited media^34^, has been previously implicated in metronidazole resistance. A single SNP present in *cbiN* distinguished a metronidazole resistant R010 isolate of *C. difficile* from a metronidazole sensitive R010 isolate isolated from the same patient^41^. In our data, *cbiN* was downregulated by 10-fold in the *D. piger* co-culture and 4-fold in *D. piger* spent media.

Another enzyme that was downregulated in the *D. piger* conditions, altronate hydrolase (*uxaA*), has potential connections with metronidazole resistance. Notably, altronate hydrolase was substantially down-regulated in *C. difficile* in the co-culture with *D. piger*, where the magnitude of this decrease in transcript abundance was the second largest in the genome (>1000-fold reduction, **Table 1, Figure 5G**). Altronate hydrolase catalyzes the dehydration of the six-carbon altronate as part of galacturonate degradation^42^. One of the 17 mutations that distinguished a metronidazole resistant NAP1 *C. difficile* strain from the metronidazole sensitive *C. difficile* reference strain occurred in the altronate hydrolase gene. The mutation in altronate hydrolase was one of three frameshift mutations in the mutant, and likely rendered altronate hydrolase non-functional^43^. Studies in *E. coli* have demonstrated that this enzyme requires iron or manganese for its catalytic activity^44^, which may explain why this gene has been observed to be strongly downregulated in iron-limited media^34^.

The mutations in *cbiN* and *uxaA* in metronidazole resistant *C. difficile* isolates suggests that the observed downregulation of *cbiN* and *uxaA* in response to iron-limitation may contribute to *C. difficile’s* increased metronidazole tolerance. These genes are potential links between iron-limitation and metronidazole tolerance, in addition to the downregulation of iron-containing oxidoreductases, hydrogenases, and ferredoxins predicted to convert metronidazole into its active form.

### Hydrogen sulfide production by D. piger leads to metal sequestration

The global changes in *C. difficile*’s gene expression profile suggest that *D. piger* created an iron-limited environment. Previous studies suggest that upregulation of *feo, fhu,* and *zupT* transporters in *C. difficile* can be induced by both zinc-starvation and iron-starvation^34, 45^ and that cross-regulation of iron and zinc has been shown in other bacteria^46^. Therefore, iron-limitation, zinc-limitation or limitation of other divalent first-row metals may contribute to the observed changes in *C. difficile* gene expression program in the presence of *D. piger*. We hypothesized that these metal limitations could have been caused by hydrogen sulfide produced by *D. piger*^47^. In *D. piger* cultures, we observed a characteristic black precipitate (ferrous sulfide) that forms when iron combines with produced hydrogen sulfide^48^. Other divalent metals can precipitate with sulfide^49, 50^ and may be precipitating in addition to ferrous sulfide in the presence of *D. piger*.

To estimate how much metal is precipitated by *D. piger* produced sulfide, we quantified the amount of sulfide in *D. piger* monoculture over time (Methods). The amount of sulfide peaked in late exponential phase at 1.4 mM (**Figure 6A**). Hydrogen sulfide is volatile and escapes during growth. Therefore, the total amount of produced hydrogen sulfide was likely higher than the measured concentration. While *C. difficile* also produces a small amount of hydrogen sulfide, the amount of hydrogen sulfide in the *C. difficile* and *D. piger* co-culture was similar to the amount in the *D. piger* monoculture (**Supplementary Figure 13A**). Based on this data, because iron was present at approximately 1.2 mM in our media and other metals were in the micromolar range (**Supplementary Figure 13B**), the produced hydrogen sulfide in the *D. piger* spent media and *D. piger* co-culture (>1.4 mM) was in excess of the total divalent metal concentration in the media. Therefore, the produced hydrogen sulfide could precipitate all divalent metals in the media, creating a metal-limited environment for *C. difficile*. This would explain the iron-limited signature in *C. difficile’s* gene expression profile. Supporting this hypothesis, iron depletion due to the precipitation with excess sulfide has been previously shown to induce Fur-regulated genes in *C. difficile*^51^.

**Figure 6:**
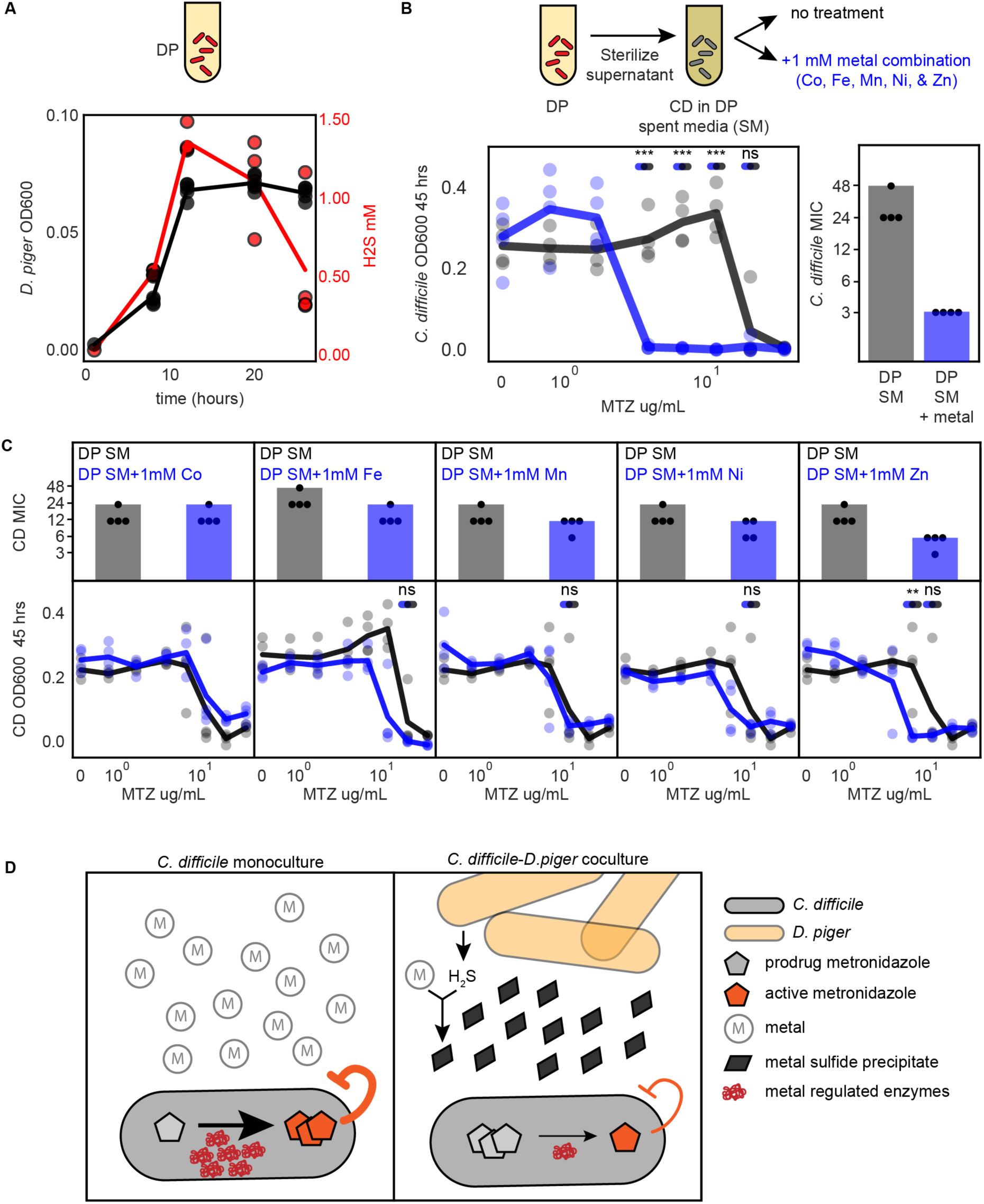
Supplementation of *D. piger* spent media with metals eliminates its protection of *C. difficile* from metronidazole. **(a)** Lineplot of absolute abundance (black) and hydrogen sulfide production (red) of *D. piger* in monoculture. Data points represent biological replicates. Each biological replicate is the average of 2 technical replicates. Line indicates average of n=4 biological replicates. **(b)** Lineplot and barplot of *C. difficile* metronidazole (MTZ) susceptibility in *D. piger* spent media (SM) with and without metal supplementation at 48 hours. Metal supplementation condition contains 1 mM of Co, Mn, Ni, Zn, and Fe. Lineplot x-axis is semi-log scale. Points represent biological replicates. Lines indicate average of n=4 biological replicates. Asterisks indicate significant difference between conditions with and without metal supplementation (*P < 0.05, **P < 0.01, ***P < 0.001) according to an unpaired t-test, “ns” indicates not significance. Statistical significance was performed at the lower of the two MICs and concentrations between the MICs of the two conditions. Barplot displays MIC of data shown in lineplot. Points represent MIC of n=4 biological replicates. Bar represents MIC determined based on average OD600 of n=4 biological replicates. **(c)** Lineplots and barplots of *C. difficile* antibiotic susceptibility in *D. piger* spent media (SM) with and without supplementation of individual metals at 48 hours. Each x-axis is semi-log scale. Points represent biological replicates. Lines indicate average of n=4 biological replicates. Barplot displays MIC of data shown in lineplot. Points represent MIC of n=4 biological replicates. Bar represents MIC determined based on average OD600 of n=4 biological replicates. Statistical significance as described in panel B. For metals with a change in MIC between the two conditions, statistical significance tested at the lower of the two MICs and any concentrations between the MICs of the two conditions. **(d)** Schematic of proposed mechanism for *D. piger* alteration of *C. difficile* metronidazole susceptibility. (Left) In monoculture, metals are available in the environment, and metal containing enzymes in *C. difficile* are expressed and reduce the prodrug metronidazole to its active form. (Right) In co-culture with *D. piger*, *D. piger* sequesters the metals via production of hydrogen sulfide, which forms metal sulfide precipitates. In response to metal limitation, the expression of metal binding proteins in *C. difficile* that reduce the conversion of metronidazole from prodrug to its active form are reduced. This in turn reduces the rate of conversion of metronidazole from its inactive to active form and increases the observed tolerance of *C. difficile* to metronidazole.

### Supplementation of D. piger spent media with combination of metals eliminates the protective effect of C. difficile from metronidazole

Based on the hypothesis that metal precipitation by hydrogen sulfide leads to an increase in *C. difficile*’s metronidazole tolerance, we tested whether removing hydrogen sulfide from *D. piger* spent media eliminates this protective effect. The majority of hydrogen sulfide was eliminated from *D. piger* spent media by purging with nitrogen gas for 15 minutes (**Supplementary Figure 14A**). We visually observed that nitrogen-purged spent media formed less ferrous sulfide precipitate than the untreated spent media (**Supplementary Figure 14B**). Consistent with the proposed mechanism, the metronidazole tolerance of *C. difficile* was reduced in the nitrogen-purged spent media compared to the untreated spent media (**Supplementary Figure 14CD**). This suggests that hydrogen sulfide contributed to the increase in *C. difficile* metronidazole tolerance via metal precipitation.

Additionally, we hypothesized that supplementing *D. piger* spent media with metals that have precipitated would also eliminate the increase in *C. difficile*’s metronidazole tolerance. We characterized *C. difficile*’s metronidazole tolerance in media supplemented with cobalt, iron, manganese, nickel, and zinc since multiple divalent metals can form sulfide precipitates^49, 50^ and limitation of multiple divalent metals can lead to similar gene expression changes in *C. difficile*^34, 45^. We introduced these metals in high concentrations (millimolar range) into the *D. piger* spent media to account for unreacted hydrogen sulfide that could precipitate the supplemented metals. *C. difficile’s* metronidazole tolerance was substantially reduced in *D. piger* spent media supplemented with the five-metal combination compared to *D. piger* spent media without this addition (16-fold decrease in MIC, **Figure 6B**). In contrast, *C. difficile*’s antibiotic tolerance was not altered in fresh media supplemented with the five-metal combination (**Supplementary Figure 15**). Supplementation of *D. piger* spent media with individual metals revealed that out of the individual metal supplementations, addition of zinc yielded the largest decrease in the protective effect (**Figure 6C**). However, none of the individual metals decreased metronidazole tolerance as substantially as the combination of metals. Overall, this data suggests that metal limitation in *D. piger* spent media caused an increase in *C. difficile* metronidazole tolerance, and metal supplementation can eliminate the protective effect of *D. piger*.

Combining our data together, we propose a biological mechanism for the effect of *D. piger* on *C. difficile*’s metronidazole tolerance (**Figure 6D**). In this proposed mechanism, hydrogen sulfide produced by *D. piger* sequesters divalent metals in the media. This in turn creates a metal limited environment for *C. difficile*, which leads to downregulation of enzymes requiring iron clusters for their activities. The down-regulated genes include iron containing enzymes that perform redox reactions that reduce metronidazole to its active form. In sum, the iron-limited environment created by *D. piger* alters the global transcriptional response in *C. difficile*, which reduces the rate of conversion of metronidazole to its active form and thus enhances the tolerance of *C. difficile* to metronidazole.

## DISCUSSION

Understanding how microbial interactions alter the antibiotic susceptibility of major human pathogens such as *C. difficile* could enable tailored antibiotic treatments informed by ecological context. These could help meet the current need for new treatments that specifically eradicate human pathogens while minimizing disruption of healthy gut microbiota and minimize the acquisition of antibiotic resistance. We investigated the contribution of inter-species interactions to the antibiotic susceptibility of a major human gut pathogen *C. difficile*. We observed two types of alterations in *C. difficile* antibiotic susceptibility: changes in *C. difficile* minimum inhibitory concentration and changes in *C. difficile* abundance at sub-inhibitory concentrations. Substantial changes in *C. difficile* tolerance were rare in our conditions, only occurring in a small fraction of characterized communities (**Supplementary Figure 16A**). However, the presence of *D. piger* substantially increased *C. difficile*’s tolerance to metronidazole, which we demonstrate is due to a reduction in bioavailable metals in the media (**Figure 5A, 6D**). By contrast, enhancements in the abundance of *C. difficile* at subMICs were more frequently observed, occurring in 52% of characterized communities (**Supplementary Figure 16B**). We demonstrate that the growth enhancement occurs via antibiotic inhibition of biotic inhibitors of *C. difficile* that display higher antibiotic sensitivity than *C. difficile* (**Figure 3C**). Our work demonstrates that pathogen growth can be altered by inter-species interactions across a wide range of antibiotic, which should be considered in the design of antibiotic treatments.

We observed ≥4-fold change in *C. difficile*’s MIC compared to monospecies in 4% of pairwise communities (**Figure 2**) and 14% of multi-species communities across both single antibiotics (**Supplementary Tables 1,2**). This is qualitatively consistent with the small fraction of inter-species interactions shown to be susceptibility modifying in other microbial communities^19–22^. The increase in *C. difficile* metronidazole tolerance by gut microbes that we observe *in vitro* may contribute to the ineffectiveness of metronidazole in treating *C. difficile* infections in the human colon. Due to the low achieved metronidazole concentration in the human colon^52^, even modest increases in *C. difficile* metronidazole tolerance could allow the pathogen to survive. Based on our results, we propose that clinical testing of *C. difficile* antibiotic susceptibility should include conditions of *C. difficile* cultured with physiologifcally relevant microbial communities. *C. difficile* could be cultured with a panel of resuspended fecal samples from multiple donors with disparate human gut microbiome compositions. Antibiotic treatments could be scored by balancing minimization of disruption of healthy gut microbiota with minimization of the variability of *C. difficile’s* MIC across donor samples.

We propose a mechanism wherein *D. piger* depletes bioavailable divalent metals (e.g., Fe and Zn) in the environment, resulting in transcriptional downregulation of enzymes in *C. difficile* requiring iron as a co-factor. These enzymes reduce metronidazole to its active form, thus protecting *C. difficile* from the action of the antibiotic (**Figure 6D**). Iron limitation has been shown to increase microbial resistance to antibiotics with proton motive force (PMF) dependent uptake due to the decrease in iron-sulfur containing complexes crucial for generating the PMF^53^. Because metronidazole passively diffuses into the cell independent of PMF^54^, iron limitation likely impacts not the initial uptake rate of metronidazole but instead the conversion rate from prodrug to activated inhibitor. Our results for metronidazole are supported by previous studies that have identified a link between diminished intracellular iron levels and *C. difficile* metronidazole resistance^39, 40^. However, we are the first to demonstrate this effect can be caused by hydrogen sulfide production produced by a constituent member of the human gut microbiome. Interestingly, heme limitation has previously been shown to sensitize *C. difficile* to metronidazole^55^, the opposite of the metronidazole protection we observed from iron-limitation, suggesting a complex relationship between heme and non-heme iron and metronidazole susceptibility.

Our identified interaction of metronidazole protection has implications beyond *D. piger* and *C. difficile*. There are multiple commensal species in addition to *D. piger* that can produce hydrogen sulfide^56^. These species may increase *C. difficile* metronidazole tolerance through a similar mechanism. Beyond *C. difficile*, metronidazole is a widely used antibiotic to treat anaerobic bacterial infections^9^, suggesting hydrogen sulfide producing bacteria could protect other pathogens from the action of metronidazole via divalent metal sequestration (**Supplementary Figure 12**).

Estimates of sulfide concentration in the colon (10^-3^ M range^57, 58^) are higher than estimates of bioavailable iron (approximately 10^-3^ M based on fecal concentrations, with 30% estimated to be in bioavailable forms^59, 60^). The excess of sulfide suggests physiological concentrations of metals may be precipitated by sulfide in the colon. However, during *C. difficile* infection, metal concentrations vary significantly as damage of epithelial cells can release heme into the environment^61^, leading to an increase in available iron. However, metal sequestration by the host immune system^62^ could simultaneously reduce available iron. The effects of sulfide on bioavailable metal in mammalian gut infections are unknown. Future studies could investigate the effect of hydrogen sulfide on *C. difficile* metronidazole susceptibility in an infection model in the murine gut. If the mechanism persists in *in vivo*, reduction of dietary protein could be considered during metronidazole treatment, as lower dietary protein has been shown to reduce hydrogen sulfide production by the gut microbiota^57, 63^.

We observed alterations in *C. difficile’s* subMIC response in approximately half of the characterized communities (**Figure 3BE**). A previous study of brewery microbiomes similarly demonstrated that antibiotic treatment reduced biotic inhibition, which in turn yielded a bloom in abundance of certain species in a pairwise and one 4-member community^21^. We demonstrate that increases in *C. difficile* abundance at subMICs can also occur in larger communities (up to a 14-membered community) and were robust across two antibiotic types (**Figure 3E**). Additionally, we showed that the magnitude of the growth enhancement depends on the degree of biotic inhibition. Finally, the presence of resistant inhibitors suppressed the growth enhancement caused by antibiotic sensitive biotic inhibitors (**Figure 3DF**, **Supplementary Figure 5AB**). Therefore, it is possible that the growth enhancement of pathogens could occur in high richness communities such as the human gut microbiome across multiple antibiotic types depending on the abundance of resistant inhibitors. Because pathogens frequently encounter subMICs during dosing regimens^25^, our data suggests narrow spectrum antibiotics or bacteriophages should be used whenever possible to specifically inhibit a pathogen and minimize disruption of antibiotic resistant biotic inhibitors to avoid pathogen growth enhancement at subMICs. Alternatively, bacterial therapeutics could be designed to introduce cocktails of resistant biotic inhibitors during treatment with broad spectrum antibiotics to suppress pathogen growth enhancement.

In sum, we demonstrate that gut microbes can substantially alter *C. difficile*’s growth response to antibiotic treatment. We provide generalizable principles to be considered in future evaluations of antibiotic treatment, including preference for narrow spectrum antibiotics and necessity of testing of pathogen MIC in the presence of physiologically relevant microbial communities.

## Methods

### Strain and culturing information

Information on strains used in this study is found in **Supplementary Table 5**. Single use glycerol stocks for each strain were created as described previously^64^. All cultures were grown in anaerobic Basal Broth (ABB, Oxoid) in an anaerobic chamber with an atmosphere of 2.5±0.5% H_2_, 15±1% CO_2_ and balance N_2_ (Coy Lab products). Starter cultures were prepared by inoculating 100 µL of single use glycerol into 5 mL of ABB. *D. piger* starter cultures were supplemented with 28mM sodium lactate (Sigma-Aldrich) and 2.7mM magnesium sulfate (Sigma-Aldrich). *E. rectale* starter cultures were supplemented with 33mM sodium acetate (Sigma-Aldrich). To ensure organisms were at similar growth stages at experimental set up, starter cultures were inoculated either 16 or 41 hours prior, depending on the organism’s growth rate (See **Supplementary Table 5**). Cultures were incubated at 37℃ with no shaking.

### Antibiotic titrations

Antibiotic susceptibility was determined following the Clinical and Laboratory Standards Institute protocol for antibiotic susceptibility of monospecies anaerobic organisms^27^, with modifications to apply the method to multispecies communities. The protocol was modified to use broth microdilution method for all conditions, because some communities contained Bacteroides species for which broth microdilution method is recommended. The broth microdilution protocol was modified to use ABB media in order to support growth of all members of the community. Lastly, the protocol was modified to determine species absolute abundance as the product of optical density and relative abundance from 16S rRNA sequencing because our conditions contained multiple species.

Metronidazole (Sigma-Aldrich M1547) and vancomycin (Sigma-Aldrich V1130) stocks of 1 mg/mL were made in water, filter sterilized with 0.2 µM filters and stored at -20C as single use aliquots. Stocks were diluted in twofold dilution series at 10X the desired concentration. Cultures were inoculated into ABB in 96 deep well plates. Monospecies were inoculated with a starting OD600 of 0.0022. Pairs were inoculated with a starting OD600 of 0.00022 for *C. difficile* and 0.00198 for other species. Multi-species communities were inoculated with a total OD600 of 0.0066 with evenness of 1. Immediately after inoculation, the 10X antibiotic dilution series was diluted 1:10 into the communities. The plates were covered with gas-permeable seals (Breathe-Easy) and incubated at 37℃ with no shaking.

After 48 hours, cultures were mixed by pipetting and 200 µL aliquots were removed for sequencing and measuring OD600. Sequencing aliquots were spun down at 1,739 *g* for 15 min after which the supernatant was removed and the pellets were stored at -80C. OD600 was measured for two dilutions of each sample and the dilution in the linear range of the instrument (Tecan Infinite Pro 200) was selected. 16S rRNA sequencing was performed to determine relative abundances as described below. Species absolute abundance was calculated by multiplying the relative abundance of each species by the community OD600. The minimum inhibitory concentration (MIC) was defined as the lowest concentration for which all concentrations greater to and equal restrict organism growth to significantly less than 0.05 OD600 as determined by a one-sample t-test.

### Determination of community composition

DNA extraction, library preparation, and sequencing were performed as described previously^4^. Briefly, cell pellets were genome extracted using the Qiagen DNEasy protocol with gram-positive lysozyme pretreatment modified for 96 well plates. Genomic DNA was normalized to 2 ng/µL in water and the 16S v3-v4 region was amplified using dual-indexed primers arrayed in 96 well plates. Samples were cleaned with DNA Clean and Concentrator kit (Zymo) and sequenced on an Illumina MiSeq.

Sequencing data was analyzed as described previously^4^. Briefly, reads were demultiplexed with Basespace FastQ Generation, paired ends were merged with PEAR v0.90^65^, and mapped to a custom database of our species using the mothur v1.40.5 command classify.seqs with the Wang method with bootstrap cutoff value of 60%^66^. Relative abundance of an organism was calculated by dividing the number of reads mapped to that organism by the total number of reads for that sample. Absolute abundance was calculated by multiplying the relative abundance by the OD600 of that sample. Samples were removed from further analysis if >1% of the reads were mapped to species not expected to be in the sample (indicating contamination).

### Generalized Lotka–Volterra Model with antibiotic perturbation

The antibiotic perturbation extension of the generalized Lotka-Volterra model^28^ is a set of *N*- coupled first-order ordinary differential equations (Equation 1):

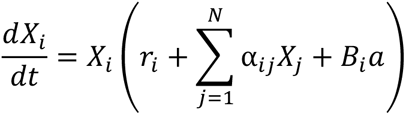

where *N* is the number of species, *X_i_* is the abundance of species *i*, *r_i_* is the basal growth rate of species *i*, *α_ij_* is the growth modification of species *i* by species *j,* and *X_j_* is the abundance of species *j*. The parameter *α_ij_* is constrained to be negative when *i* = *j*, representing intra-species competition. In the antibiotic term, *B_i_* is the sensitivity of species *i* to antibiotic, and a is the concentration of the antibiotic. We modify the model to make the antibiotic concentration modified constant over time.

Values for growth rates parameters *r_i_* and interaction parameters *a_ij_* come from our previous work^4^. Values for antibiotic sensitivity parameters *B_i_* were inferred in this study from time-series measurements of monospecies antibiotic titrations using a custom python script. The equation used to infer *B_i_* from monospecies data was Equation 1, with *α_ij_* = 0 for all j!=i. The minimize function of the Scipy optimize package was used to determine the optimal B*i* for each species x_*i*_, that resulted in the lowest cost between the antibiotic gLV model and the time-series measurements across all the antibiotic concentrations in the titration. The antibiotic concentrations were scaled so that the concentrations range between 0 and 1. The cost in the optimization was computed by simulating the species abundance in each antibiotic condition with an ODE solver and summing the mean-squared error between the abundance of the species in the simulation and the abundance of the species in the data at each timepoint in each antibiotic concentration.

### Null computational models

Null model 1 (“gLV”) is the generalized Lotka-Volterra Model. This is the same as the generalized Lotka-Volterra Model with antibiotic perturbation (Equation 1) without the antibiotic term. This can also be thought of as the generalized Lotka-Volterra with antibiotic perturbation with all species antibiotic susceptibility set to zero (not susceptible). Null model 1 is Equation 1 with *B_i_*=0 for all species.

Null model 2 (“gLV + shuffled antib. suscept.”) is the generalized Lotka-Volterra Model with antibiotic perturbation (Equation 1) with a shuffled set of antibiotic susceptibility parameters. For metronidazole conditions, the 14 metronidazole susceptibility parameters were shuffled and for the vancomycin conditions, the 14 vancomycin susceptibility parameters were shuffled (the shuffled *B_i_* for species i is equal to the unshuffled *B* of any one of the 14 species). The accuracy of Null model 2 was averaged across 1000 sets of shuffled parameters. The distance between the shuffled B for *C. difficile* and the unshuffled B for *C. difficile* was calculated as the absolute abundance of the difference between shuffled *B*_CD_ and unshuffled *B*_CD_.

Null model 3 (“gLV + monosp. Antib. Susceptibility, aijs=0”) is the generalized Lotka-Volterra Model with antibiotic perturbation (Equation 1) with no interspecies interactions. Null model 3 is Equation 1 with *α_ij_* = 0 for all j!=i.

### Spent media preparation

*D. piger* starter culture was incubated for 48 hours. The culture was then diluted into ABB to an OD600 of 0.0022. The cultures were incubated at 37℃ with no shaking. Unless otherwise indicated, cultures were incubated for 12 hours. After incubation, cultures were spun down at 1,739 *g* for 15 min. Supernatant was removed and filter sterilized with Steriflip 0.2 µM filters.

### Metronidazole incubation experiment

*D. piger* spent media was prepared following the procedure described above. An aliquot of metronidazole was incubated in *D. piger* spent media or rich media (ABB) at 37C. After 12 hours of incubation, the incubated metronidazole was diluted into a *C. difficile* culture in fresh media (ABB) following the antibiotic titration method described above.

### Iron-sulfur cluster prediction

The protein FASTA sequence from the *C. difficile* R20291 reference genome (GenBank assembly accession GCF_000027105.1) was run on the Metal Predator web-server^29^.

### Metal quantification and supplementation

The amount of metal in fresh media (Anaerobic Basal Broth) was analyzed via Inductively Coupled Plasma Mass Spectroscopy by the Wisconsin State Lab of Hygiene.

In metal supplementation experiment, fresh media (Anaerobic Basal Broth) was supplemented with iron (II) sulfate (Alfa Aesar), manganese (II) sulfate monohydrate (Sigma-Aldrich), nickel sulfate (II) hexahydrate (Sigma-Aldrich), zinc (II) sulfate heptahydrate (Alfa Aesar), and/or cobalt (II) chloride hexahydrate (Sigma-Aldrich). Metal compounds were prepared as 100x stocks in water and filter sterilized with 0.2 µM filters.

### Hydrogen sulfide quantification

Hydrogen sulfide was quantified using the Cline Assay. Immediately before performing the assay, sodium sulfide (Alfa Aesar) was added to nitrogen purged water into a sealed vial. The stock was then diluted into 1% zinc acetate to 1mM and further diluted into 1% zinc acetate to desired concentrations for a standard curve. Samples were removed from the anaerobic chamber and immediately diluted 5-fold into 1% zinc acetate. Cline reagent was prepared in advance (1.6 g N,N-dimethyl-p-phenylenediamine sulfate (Acros Organics), 2.4g iron (III) chloride (Spectrum Chemical), 50mL concentrated hydrochloric acid, 50 mL water, stored in the dark) of which 3 µL was added to the standards and samples. Samples were mixed by pipetting and incubated for 20 minutes in the dark before measuring absorbance at 670nm. If the samples contained cells, after incubation the samples were spun down at 1,739 *g* for 10 min and the supernatant was transferred to a new plate to measure OD670.

### Transcriptomics

*D. piger* spent media was prepared following the procedure described above. *C. difficile* and *D. piger* starter cultures were incubated for 48 hours. *C. difficile* monoculture and CD-DP co-culture conditions were inoculated from starter cultures into 96 deep well plates. For monoculture conditions, *C. difficile* was inoculated to an OD600 of 0.0022. For CD-DP co-culture, *C. difficile* was inoculated to an OD600 of 0.0022 and *D. piger* was inoculated to an OD600 of 0.0198. Immediately after inoculation, antibiotics were added as a 1:10 dilution of 10X antibiotic stocks. The plates were covered with gas-permeable seals (BreathEasy) and were incubated at 37℃ with no shaking. After 14 hours, 800 µL of RNAprotect (Qiagen) was added to 400 µL of culture after which the cultures were mixed by pipetting and then incubated for 5 min at room temperature. Cultures were then centrifuged at room temperature for 10 minutes at 1,739 *g* and supernatant was carefully removed. Pellets were stored at -80C.

RNA was extracted using acidic phenol bead-beating method. Pellets were resuspended in 500 µL 2X Buffer B (200mM sodium chloride, 20mM ethylenediaminetetraacetic acid) and transferred to 2 mL microcentrifuge tubes containing 500 µL Phenol:Chloroform:IAA (125:24:1, pH 4.5) and 210 µL 20% sodium dodecyl sulfate and were bead-beated with acid washed beads (Sigma G1277) for 3 minutes. All solutions were RNAse-free. Samples were centrifuged at 4C for 5 minutes at 8000rpm, and 600 µL of the upper aqueous phase was added to 60 µL cold isopropanol and chilled on ice for 5 minutes before freezing for 5 minutes at -80C. Samples were centrifuged at 4C for 15 minutes at 18200g, the supernatant was decanted, and the pellet was washed with cold 100% ethanol. The pellets were dried in a biosafety cabinet for 10 minutes and then resuspended in 100 µL RNAse-free water. Samples were purified using RNeasy Mini Kit (Qiagen) and genomic DNA was removed using RNAse-Free DNase Set (Qiagen). Two replicates of each condition were sent to GENEWIZ (NJ, USA) for sequencing. GENEWIZ depleted rRNA with Ribozero rRNA Removal Kit (Illumina) before cDNA library preparation using NEBNext Ultra RNA Library Prep (NEB). GENEWIZ sequenced the libraries on Illumina HiSeq. Data was de-multiplexed using Illumina’s bcl2fastq 2.17 software, where one mismatch was allowed for index sequence identification.

The data was quality checked using FastQC. The BBDuk, BBSplit, and BBMap tools from BBTools suite were used to trim adapters, deplete rRNA, and map the remaining mRNA reads to the reference genomes (*C. difficile:* GenBank assembly accession GCF_000027105.1. *D. piger:* Genbank assembly accession GCA_000156375.1). FeatureCounts was used to map reads to features on the genome. RPKM values were computed using a custom python script. The DESeq2 Bioconductor library v4.0.3 was used in R v4.0.4 to quantify differential gene expression using a negative binomial generalized linear models with apeglm shrinkage estimator. When calculating RPKM of *C. difficile* genes in the *C. difficile*-*D. piger* co-culture, the “reads mapped” in the denominator was the number of reads mapped to the *C. difficile* genome. Similarly, when quantifying differential gene expression for *C. difficile* genes in the *C. difficile*-*D. piger* co-culture, only reads mapped to the *C. difficile* genome were provided to DeSeq2.

### Gene set enrichment analysis

Gene set enrichment analysis was performed using the GSEA method of the ClusterProfiler R package (v4.2.2)^67^. KEGG modules for *C. difficile* R20291 (KEGG T number T00998) were used as gene sets and were supplied as a user defined annotation with the TERM2GENE field. The analysis was run with the log2FCs calculated by DeSeq2. The p value cutoff used was 0.05 and the minimum gene set size used was 3.

## ACKNOWLEDGEMENTS

The authors would like to thank Dr. Tricia Kiley for helpful discussions. Research was sponsored by the National Institutes of Health and was accomplished under grant number R35GM124774. S.H. was supported in part by the National Institute of General Medical Sciences of the National Institutes of Health under Award Number T32GM008349.

## AUTHOR CONTRIBUTIONS

O.S.V. and S.H. conceived the study. S.H. carried out the experiments and performed computational modeling and analysis. S.H. and O.S.V. analyzed and interpreted the data and wrote the paper. O.S.V. secured funding.

