## Supplementary Figures for "Microbial interactions impact the growth response of *Clostridioides difficile* to antibiotics"

**A**

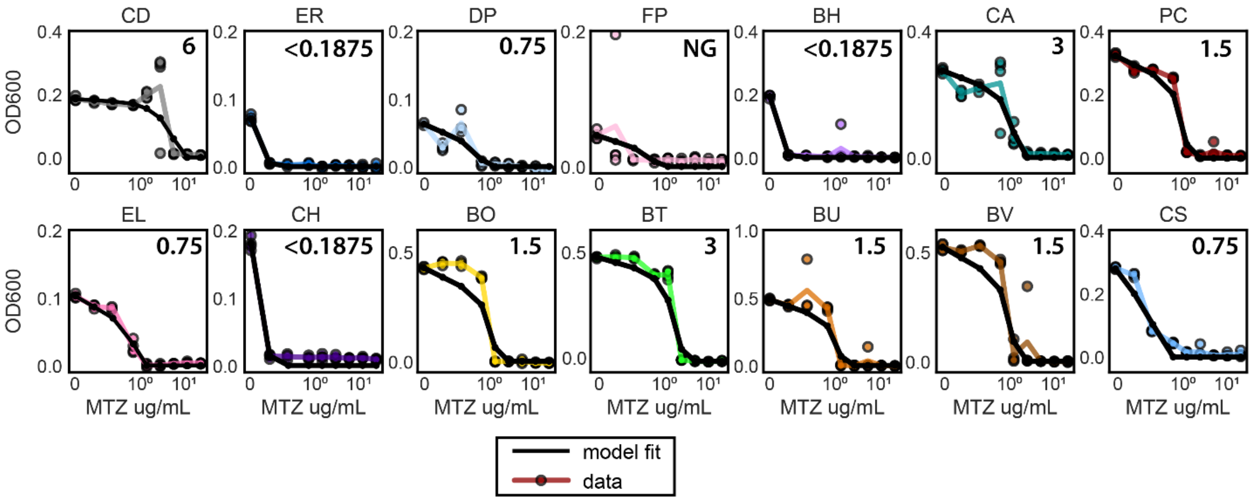


**B**

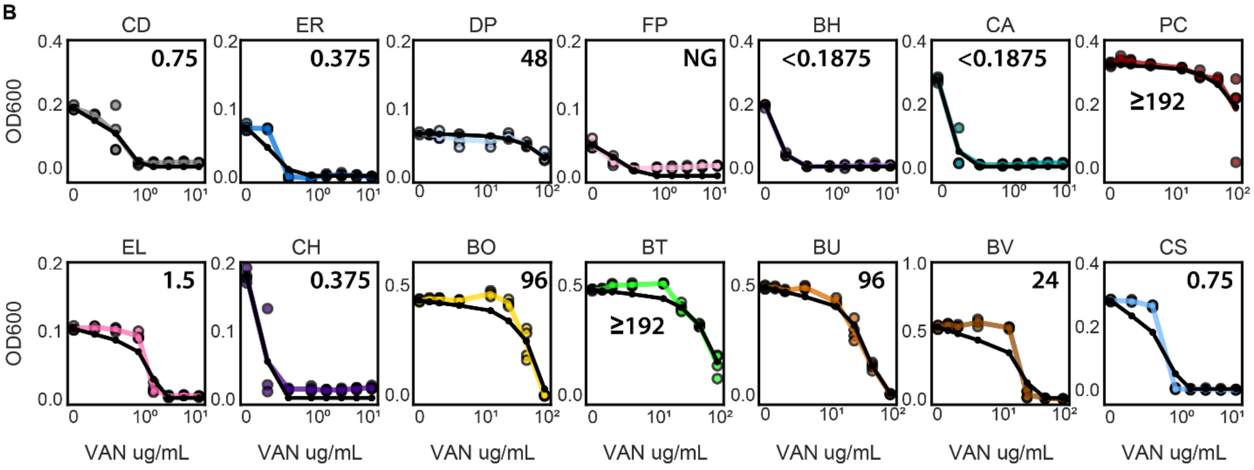


**C**

| Species | Scaled B (MTZ) | Scaled B (VAN) |
|---|---|---|
| CD | -0.9 | -4.7 |
| ER | -30 | -6.8 |
| DP | -5.2 | -0.1 |
| FP | -4.6 | -4.5 |
| BH | -35 | -14 |
| CA | -3.7 | -17 |
| PC | -2.7 | -0.1 |
| EL | -4.6 | -1.2 |
| CH | -25 | -9.6 |
| BO | -5.9 | -0.4 |
| BT | -3.4 | -0.4 |
| BU | -6.4 | -0.7 |
| BV | -4.8 | -1.2 |
| CS | -8.5 | -2.7 |

**Supplementary Figure 1: Metronidazole and Vancomycin tolerance of *C. difficile* and gut microbes in monoculture. (a) (b)** Lineplots of monospecies OD600 across antibiotic concentrations for metronidazole (MTZ) and vancomycin (VAN) at 48 hours. Each x-axis is semi-log scale. Colored data points indicate biological replicates. Colored lines indicate the average of n=2 to n=4 biological replicates. Bold number indicates MIC of data. Black lines indicate model fit. **(c)** Table of scaled antibiotic susceptibility parameter B for each species.

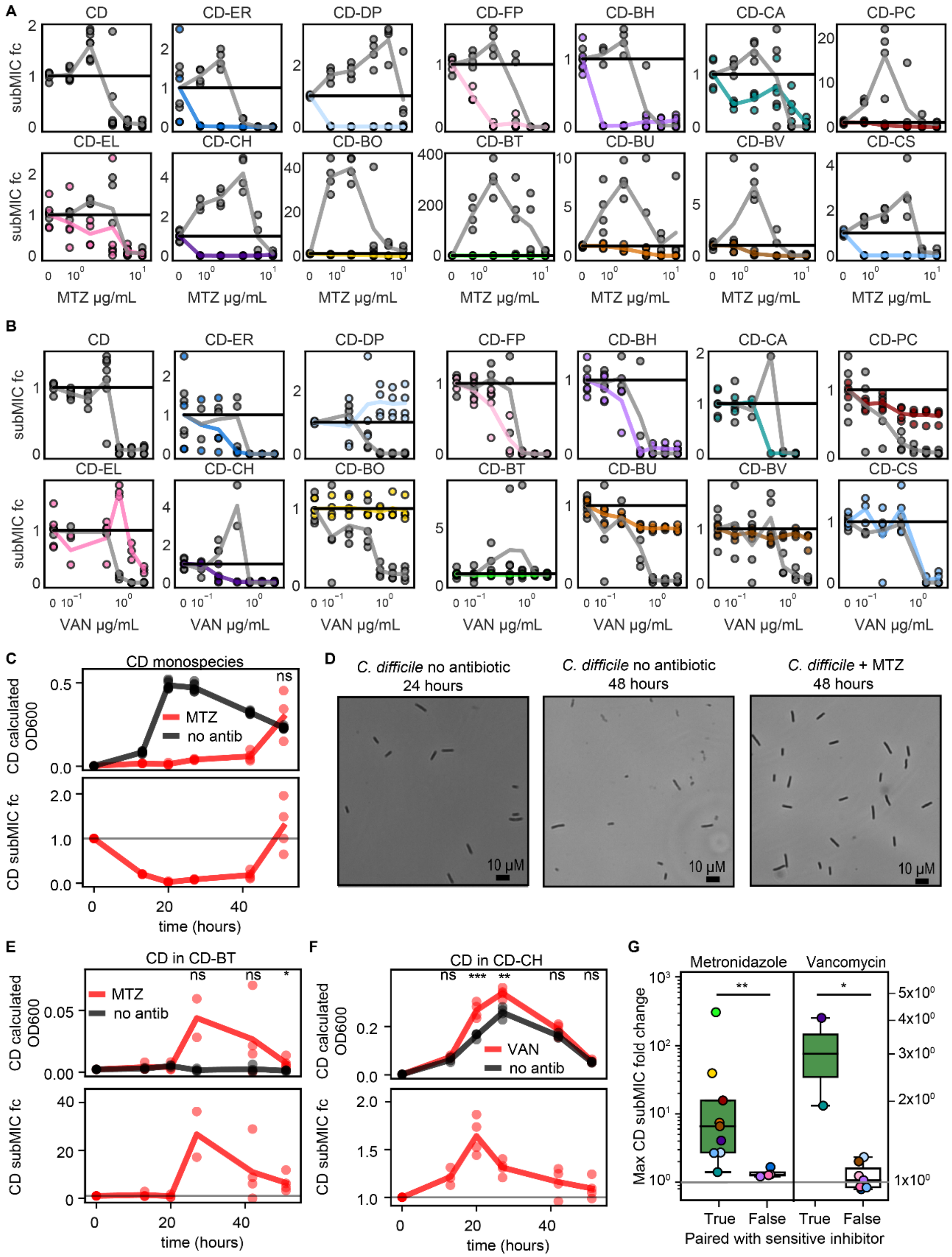


**Supplementary Figure 2: SubMIC fold change for *C. difficile*-gut microbe pairs. (a) (b)** Lineplots of subMIC fold change for pairs in metronidazole (MTZ) and vancomycin (VAN) at 48 hours. SubMIC fold change is calculated as the species absolute abundance at the subMIC divided by the species absolute abundance in the no antibiotic condition. Black horizontal line at y=1 indicates no change in growth compared to no antibiotic condition**.** Each x-axis is semi-log scale. Points indicate biological replicates. Line represents mean of n=3 to n=8 biological replicates in panel A and n=1 to n=8 biological replicates in panel

B . **(c)** Lineplots of *C. difficile* monospecies over time. (Top) *C. difficile* OD600 at 0 μg/mL and 1.5 μg/mL metronidazole. (Bottom) *C. difficile* subMIC fold change μg/mL metronidazole. Gray horizontal line at y=1 indicates no change in growth compared to no antibiotic condition. SubMIC fold change calculated as in panels AB. Points represent biological replicates. Line represents average of n=4 biological replicates. Statistical significance analyzed for all timepoints where *C. difficile* OD600 was greater in the presence of antibiotic than the absence of antibiotic ("ns" P>0.05, according to an unpaired t-test). **(d)** Microscopy images of select *C. difficile* conditions from panel C: no antibiotic at 24 and 48 hours and 1.5 μg/mL metronidazole at 48 hours. Exponential phase *C. difficile* cells in presence of subMIC of metronidazole do not show morphological difference from exponential phase *C. difficile* cells grown in absence of antibiotic. **(e)** Lineplots of *C. difficile* in pair with *Bacteroides thetaiotaomicron* over time. (Top) *C. difficile* OD600 in the presence of 0 μg/mL or 1.5 μg/mL metronidazole. (Bottom) *C. difficile* subMIC fold change in the presence of 1.5 μg/mL metronidazole. Gray horizontal line at y=1 indicates no change in growth compared to no antibiotic condition. SubMIC fold change calculated as in panels AB. Points represent biological replicates. Line represents average of n=4 biological replicates. Statistical significance analyzed for all timepoints where *C. difficile* OD600 was greater in the presence of antibiotic than the absence of antibiotic ("ns" P>0.05, according to an unpaired t-test). **(f)** Lineplots of *C. difficile* in pair with *Clostridium hiranonis* over time. (Top) *C. difficile* OD600 in the presence of 0 μg/mL or 0.1875 μg/mL vancomycin. (Bottom) *C. difficile* subMIC fold change in the presence of 0.1875 μg/mL vancomycin. Gray horizontal line at y=1 indicates no change in growth compared to no antibiotic condition. SubMIC fold change calculated as in panels AB. Points represent biological replicates. Line represents average of n=4 biological replicates. Statistical significance analyzed for all timepoints where *C. difficile* OD600 was greater in the presence of antibiotic than the absence of antibiotic (*P < 0.05, **P < 0.01, ***P < 0.001, ns P>0.05, according to an unpaired t-test). **(g)** Box plot of maximum subMIC fold change for *C. difficile* in pairs at 48 hours. The maximum subMIC fold change is the maximum of the average subMIC fold change at all subMICs. The average subMIC fold change was calculated for each subMIC concentration by taking the average *C. difficile* absolute abundance at the subMIC divided by the average *C. difficile* absolute abundance in the no antibiotic condition, where the average is of n=1 to n=4 biological replicates. *C. difficile* did not grow at any concentrations of vancomycin in CD-BO, CD-BT, CD-BU, and CD-PC pairs so was not included in the analysis. Each data point represents one pair. Gray horizontal line at y=1 indicates no change in growth compared to no antibiotic condition. Species are categorized as sensitive inhibitors if 1) the absolute abundance of *C. difficile* in pair with that species was significantly lower than the absolute abundance of *C. difficile* in monospecies, in the absence of antibiotics, as determined by an unpaired t-test and 2) if the species monospecies MIC was less than the monospecies MIC of *C. difficile*. Asterisks indicate significant difference (*P < 0.05, **P < 0.01, ***P < 0.001) according to a one-sided Mann–Whitney U test.

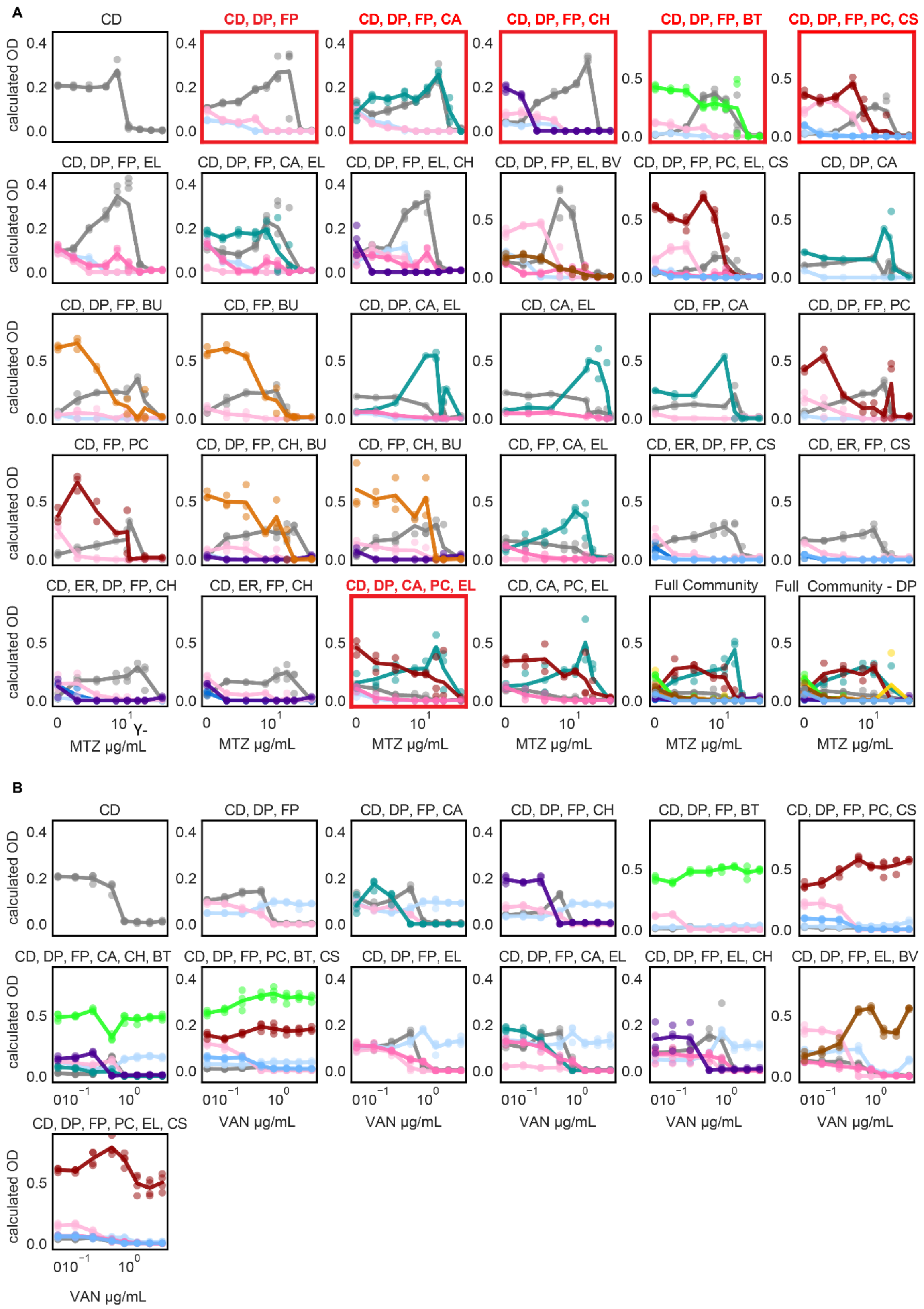


**Supplementary Figure 3: Metronidazole and vancomycin tolerance of multispecies communities. (a) (b)** Lineplots of absolute abundance across A metronidazole (MTZ) and B vancomycin (VAN)

concentrations at 48 hours. Each x-axis is semi-log scale. Y-axis is calculated OD600 (OD600 multiplied by relative abundance from 16S sequencing). Data points indicate biological replicates. Lines indicate the average of n=1 to n=4 biological replicates in panel A and n=3 to n=4 biological replicates in panel B. Color indicates species, see Figure 1C. Red borders indicate communities with ≥4-fold change in *C. difficile* MIC (see Supplementary Table 1).

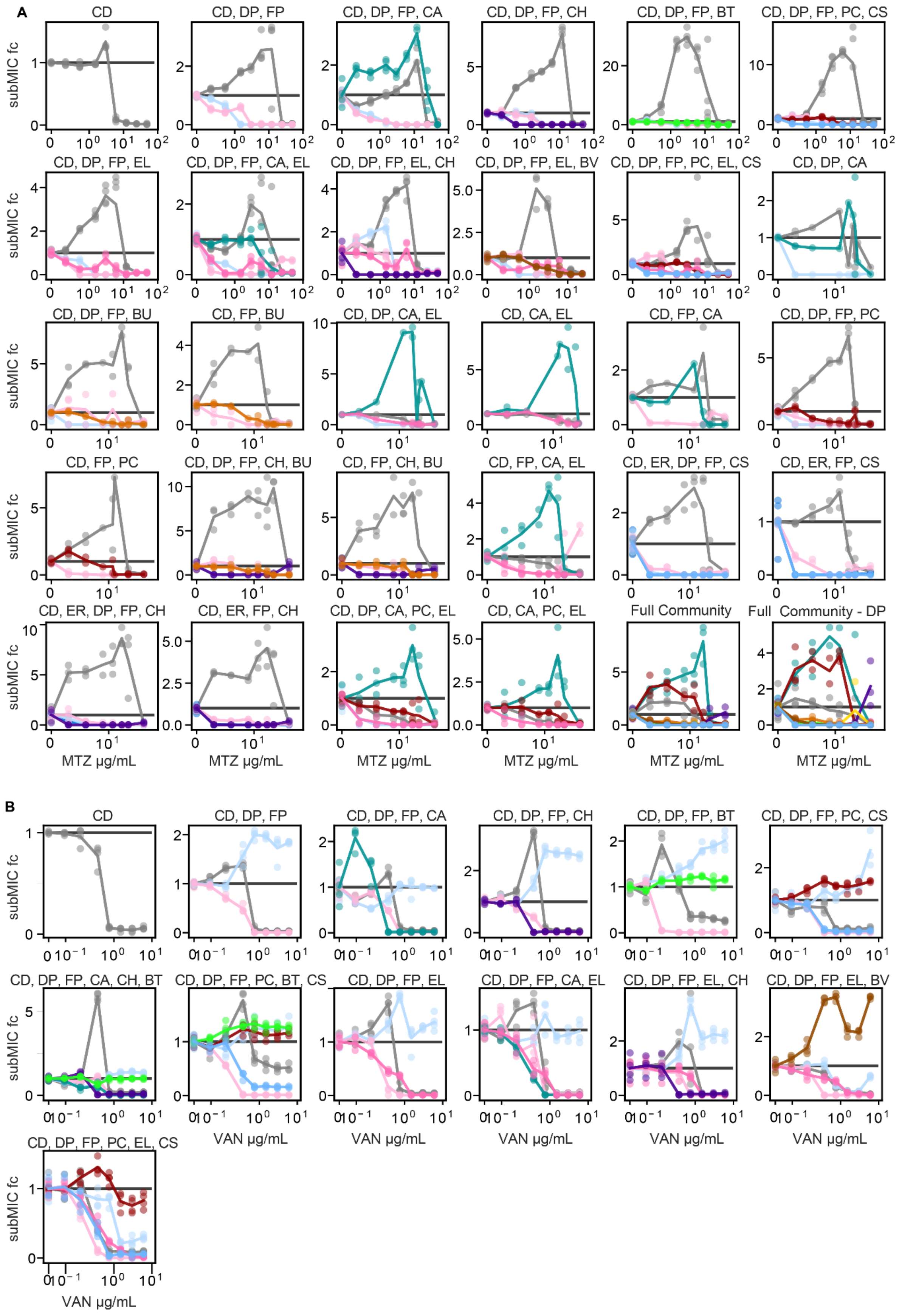
A
CD
CD, DP, FP
CD, DP, FP, CA
CD, DP, FP, CH
CD, DP, FP, BT
CD, DP, FP, PC, CS
CD, DP, FP, EL
CD, DP, FP, CA, EL
CD, DP, FP, EL, CH
CD, DP, FP, EL, BV
CD, DP, FP, PC, EL, CS
CD, DP, CA
CD, DP, FP, BU
CD, FP, BU
CD, DP, CA, EL
CD, CA, EL
CD, FP, CA
CD, DP, FP, PC
CD, FP, PC
CD, DP, FP, CH, BU
CD, FP, CH, BU
CD, FP, CA, EL
CD, ER, DP, FP, CS
CD, ER, FP, CS
CD, ER, DP, FP, CH
CD, ER, FP, CH
CD, DP, CA, PC, EL
CD, CA, PC, EL
Full Community
Full Community - DP
subMIC fc
MTZ μg/mL
B
CD
CD, DP, FP
CD, DP, FP, CA
CD, DP, FP, CH
CD, DP, FP, BT
CD, DP, FP, PC, CS
CD, DP, FP, CA, CH, BT
CD, DP, FP, PC, BT, CS
CD, DP, FP, EL
CD, DP, FP, CA, EL
CD, DP, FP, EL, CH
CD, DP, FP, EL, BV
CD, DP, FP, PC, EL, CS
subMIC fc
VAN μg/mL

**Supplementary Figure 4: SubMIC fold change for *C. difficile* in multispecies communities. (a) (b)** Lineplots of subMIC fold change in metronidazole (MTZ) and vancomycin (VAN) at 48 hours. SubMIC fold change is calculated as the species absolute abundance at the subMIC divided by the species absolute abundance in the no antibiotic condition. Black horizontal line at y=1 indicates no change in growth compared to no antibiotic condition**.** Each x-axis is semi-log scale. Data points indicate biological replicates. Lines indicate the average of n=1 to n=4 biological replicates in panel A and n=3 to n=4 biological replicates in panel B. Color indicates species, see Figure 1C.

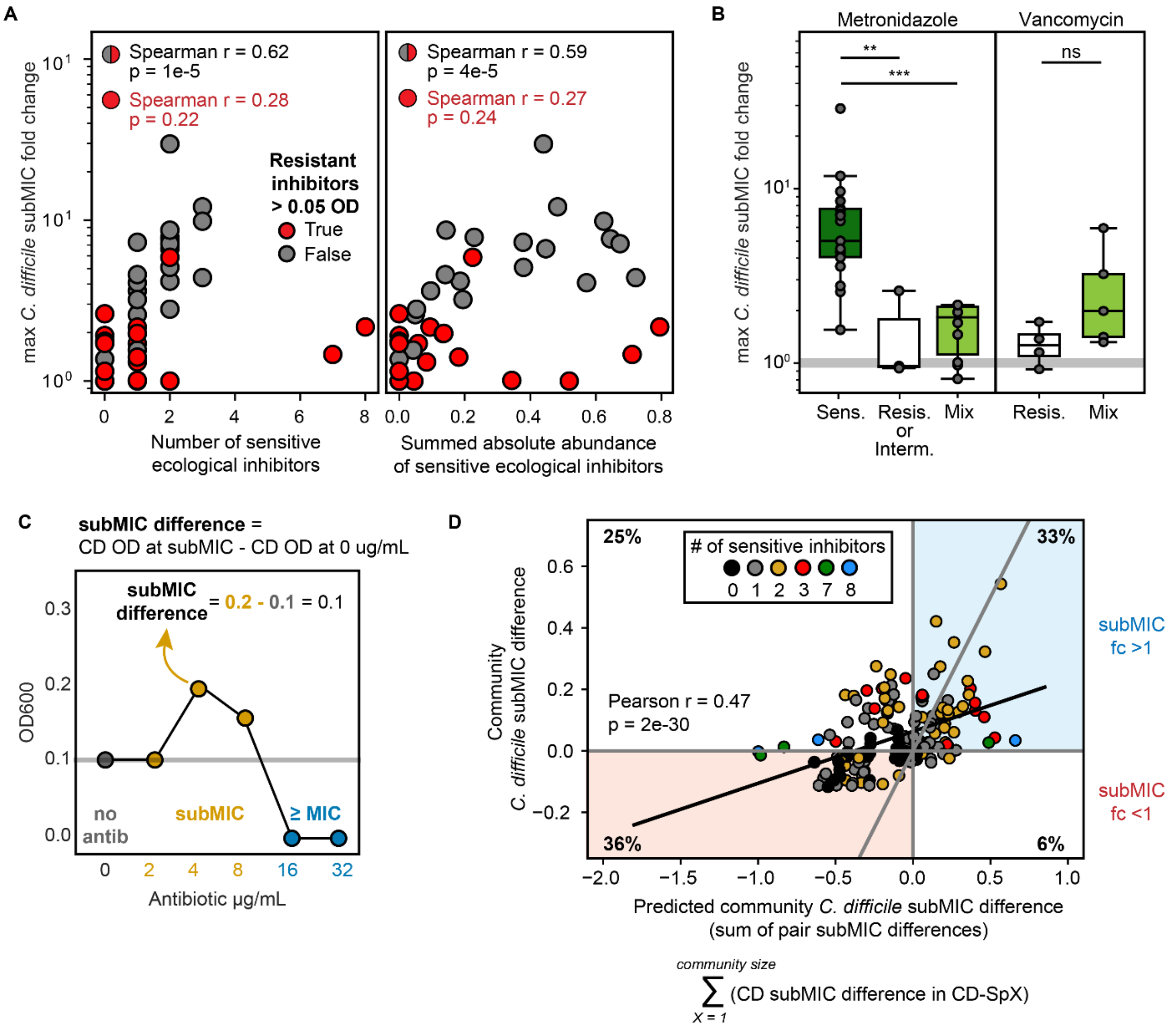


**Supplementary Figure 5: Analysis of subMIC fold change for *C. difficile* in multispecies communities. (a)** Scatterplots of maximum *C. difficile* subMIC fold change in multispecies communities at 48 hours as a function of number (left) or amount (right) of sensitive biotic inhibitors for both metronidazole and vancomycin. X-axis is number of sensitive biotic inhibitors in community at community inoculation (left) or summed absolute abundance of sensitive biotic inhibitors in community at 48 hours in the no antibiotic condition (right). The average subMIC fold change was calculated for each subMIC concentration by taking the average *C. difficile* absolute abundance at the subMIC divided by the average *C. difficile* absolute abundance in the no antibiotic condition, average of n=1 to n=4 biological replicates. The maximum subMIC fold change (y-axis) is the maximum of the average subMIC fold change at all subMICs. The sensitive and resistant inhibitors are determined as in Fig3F. Red points indicate communities where sum of absolute abundance of resistant biotic inhibitors at 48 hours was greater than 0.05 OD600. Spearman correlation annotated for all data points (black) and for only communities with resistant inhibitors > 0.05 OD600 (red). **(b)** Box plot of maximum subMIC fold change for *C. difficile* in multispecies communities. The maximum subMIC fold change (y-axis) is calculated as in panel A. Each data point represents one community. Community type (x-axis) determined as in Figure 3F. Gray horizontal line at y=1 indicates no change in growth compared to no antibiotic condition. Asterisks indicate significant difference (*P < 0.05, **P < 0.01, ***P < 0.001) according to a one-sided Mann–Whitney U test. **(c)** Schematic of subMIC difference metric. The x-axis is semi-log scale. Gray horizontal line at y=1 indicates no change in growth compared to no antibiotic condition. **(d)** Scatterplots comparing predicted value of *C. difficile* subMIC difference in community (x-axis) with average experimental value of *C. difficile* subMIC difference in community (y-axis). Predicted value was calculated by summing the *C. difficile* subMIC differences of each CD-SpX pair for all SpX in the community. SubMIC differences greater than zero have subMIC fold changes greater than one. SubMIC differences less than zero have subMIC fold changes less than one. Each point represents one community at one concentration of metronidazole or vancomycin. Number of sensitive inhibitors (color) determined as in Figure3F. Black line indicates best fit linear regression for all data points. Gray y=x line represents perfect prediction of community subMIC difference. Annotated pearson correlation values are from linear regression of all data points. Annotated percentages indicate the percentage of total data points in a given quadrant.

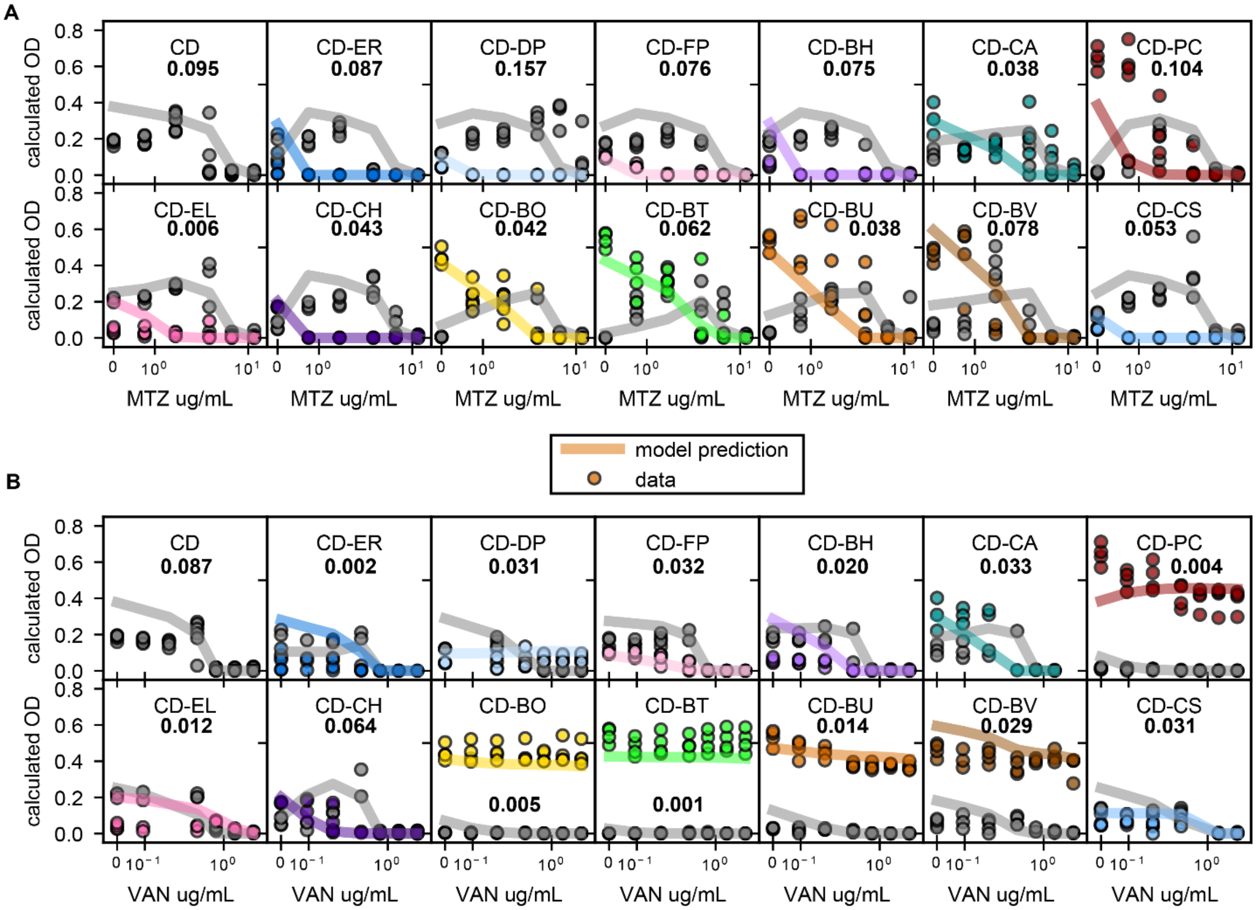

**Supplementary Figure 6: Antibiotic gLV model prediction of pairs data**. **(a) (b)** Lineplots of model prediction of absolute abundance of *C. difficile* and gut microbe pairs at 48 hours in presence of A metronidazole (MTZ) or B vancomycin (VAN). Each x-axis is semi-log scale. Y-axis is calculated OD600 (OD600 multiplied by relative abundance from 16S sequencing). Points indicate experimental data. Lines indicate model simulations. Color indicates species, see Figure 1C. Bold number is sum of squared errors for *C. difficile* (square of difference between model OD600 and average experimental OD600, summed across all concentrations).

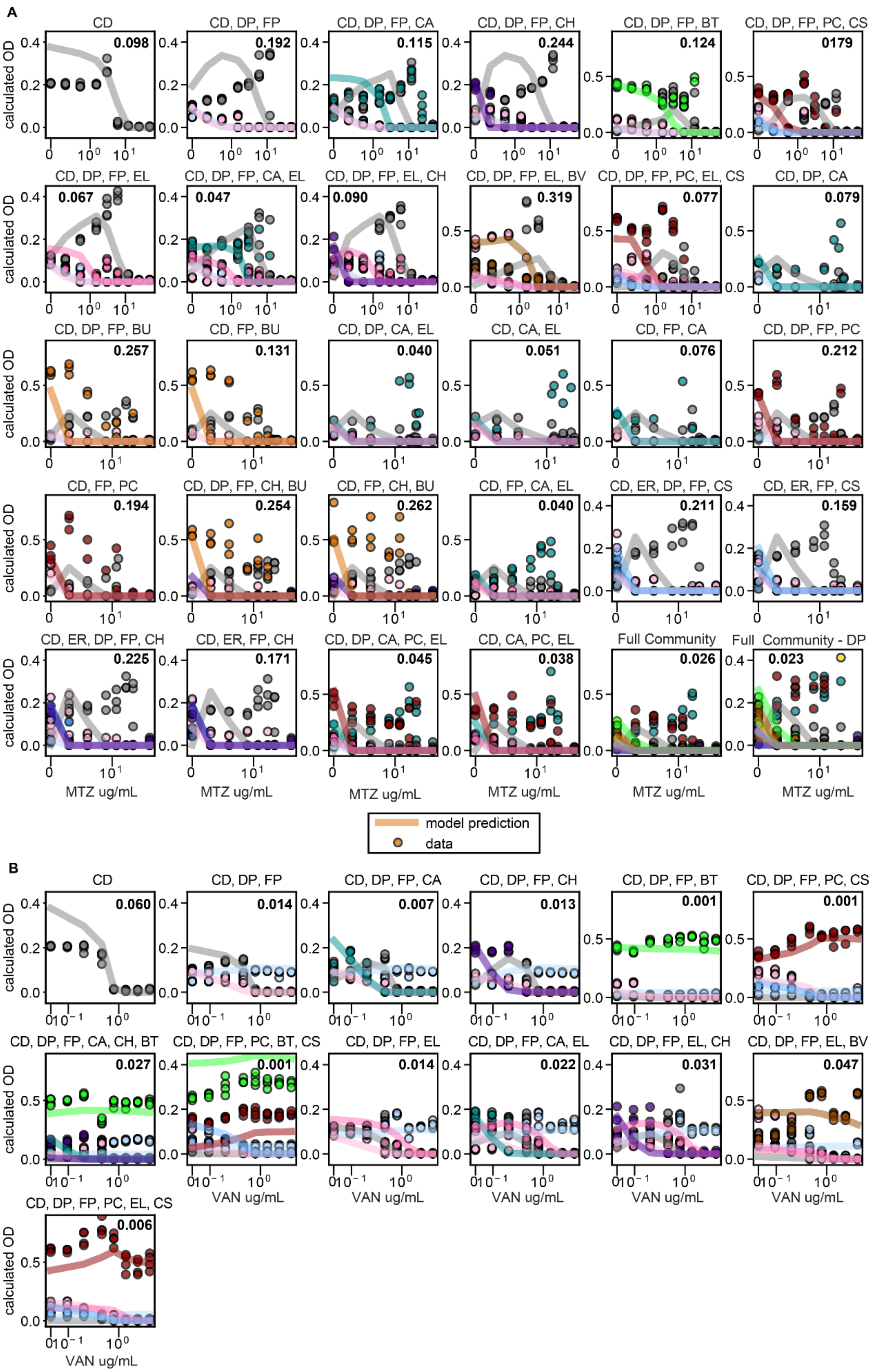
A
CD
0.098
CD, DP, FP
0.192
CD, DP, FP, CA
0.115
CD, DP, FP, CH
0.244
CD, DP, FP, BT
0.124
CD, DP, FP, PC, CS
0179
CD, DP, FP, EL
0.067
CD, DP, FP, CA, EL
0.047
CD, DP, FP, EL, CH
0.090
CD, DP, FP, EL, BV
0.319
CD, DP, FP, PC, EL, CS
0.077
CD, DP, CA
0.079
CD, DP, FP, BU
0.257
CD, FP, BU
0.131
CD, DP, CA, EL
0.040
CD, CA, EL
0.051
CD, FP, CA
0.076
CD, DP, FP, PC
0.212
CD, FP, PC
0.194
CD, DP, FP, CH, BU
0.254
CD, FP, CH, BU
0.262
CD, FP, CA, EL
0.040
CD, ER, DP, FP, CS
0.211
CD, ER, FP, CS
0.159
CD, ER, DP, FP, CH
0.225
CD, ER, FP, CH
0.171
CD, DP, CA, PC, EL
0.045
CD, CA, PC, EL
0.038
Full Community
0.026
Full Community - DP
0.023
calculated OD
MTZ ug/mL
model prediction
data
B
CD
0.060
CD, DP, FP
0.014
CD, DP, FP, CA
0.007
CD, DP, FP, CH
0.013
CD, DP, FP, BT
0.001
CD, DP, FP, PC, CS
0.001
CD, DP, FP, CA, CH, BT
0.027
CD, DP, FP, PC, BT, CS
0.001
CD, DP, FP, EL
0.014
CD, DP, FP, CA, EL
0.022
CD, DP, FP, EL, CH
0.031
CD, DP, FP, EL, BV
0.047
CD, DP, FP, PC, EL, CS
0.006
calculated OD
VAN ug/mL

**Supplementary Figure 7: Antibiotic gLV model prediction of multispecies community data. (a) (b)** Lineplots of model prediction of absolute abundance of multispecies communities at 48 hours in presence of A metronidazole (MTZ) or B vancomycin (VAN). Each x-axis is semi-log scale. Y-axis is calculated OD600 (OD600 multiplied by relative abundance from 16S sequencing). Points indicate experimental data. Lines indicate model simulations. Color indicates species, see Figure 1C. Bold number is sum of squared errors for *C. difficile* (square of difference between model OD600 and average experimental OD600, summed across all concentrations).

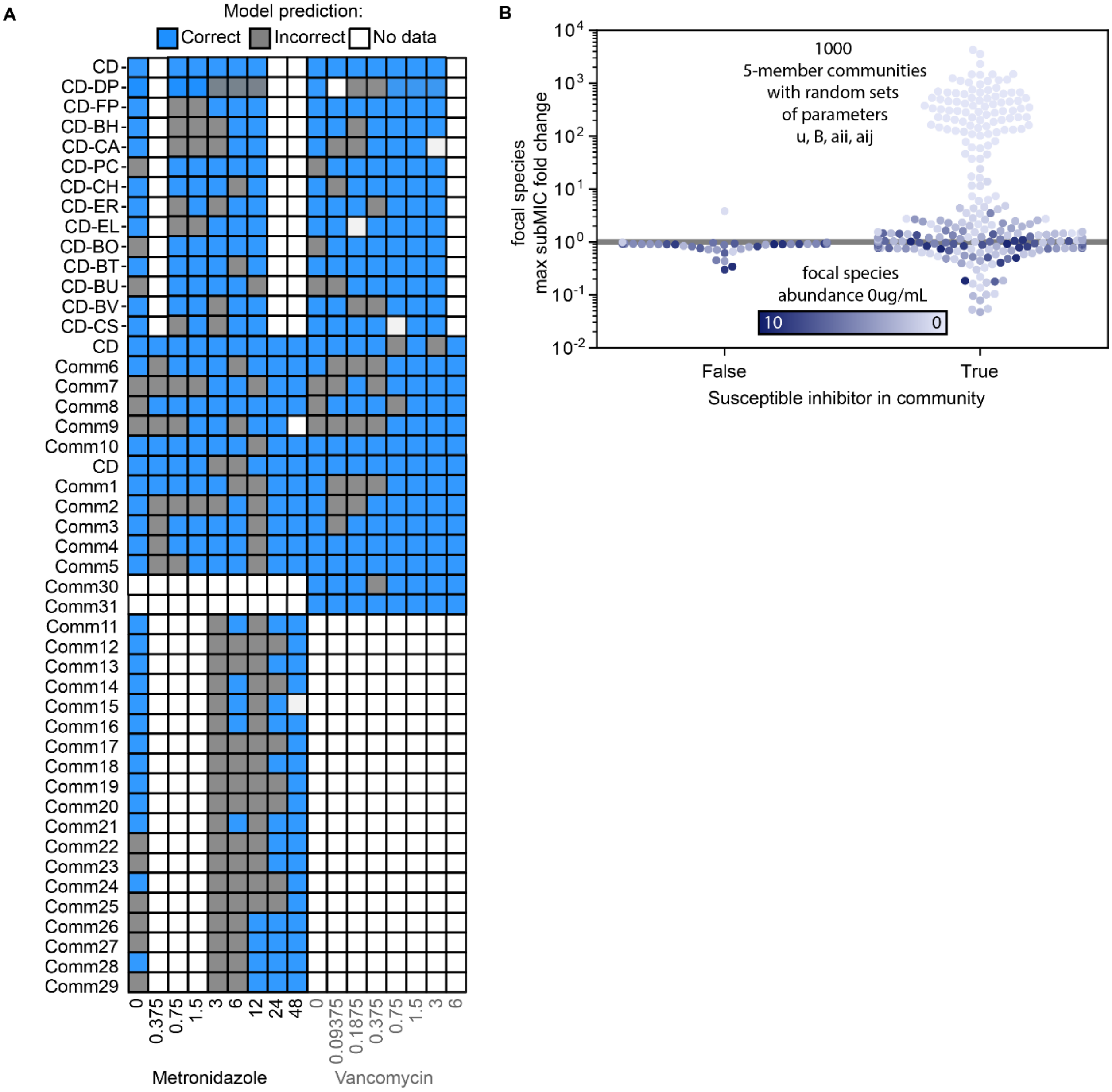

**Supplementary Figure 8: Modelling subMIC fold change. (a)** Antibiotic gLV model prediction of pairs and multispecies community data at each antibiotic concentration. These predictions are what is summarized in the blue “gLV + monospecies antib susceptibility” bar in Figure4B. **(b)** Simulated max subMIC fold change for a focal species cocultured with a four other species for 1000 random parameter sets. All parameters were randomized between a set of bounds. Bounds: aji (-1.25, 1.25), growth rates (0,1), intraspecies interactions (-1.25, 0), and antibiotic susceptibility (-6,0). Color of datapoint indicates abundance of focal species in the community under no antibiotics (where light colored data points indicate inhibition of focal species). Community is classified as containing a susceptible inhibitor if for any non-focal species j, $B_j$ < -0.1 and $a_{focal,j}$ < -0.1.  Gray horizontal line at y=1 indicates no change in growth compared to no antibiotic condition

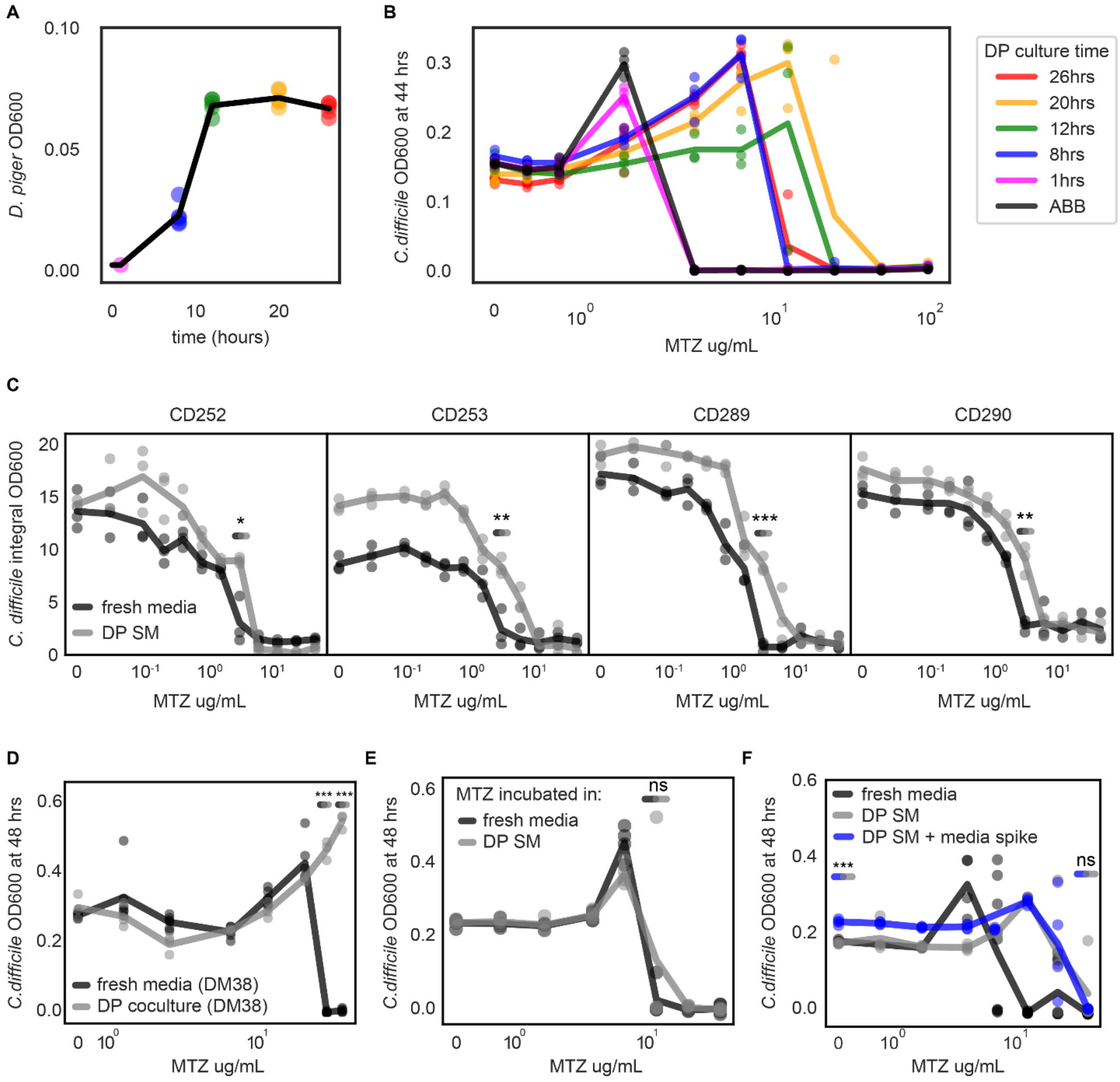

**Supplementary Figure 9: Robust protection of *C. difficile* from metronidazole by spent media of *D. piger.*** **(a)** Line plot of growth curve *D. piger*. Colored points indicate time at which five *D. piger* spent media samples were harvested. Points represent biological replicates. Line represents mean of n=5 biological replicates. **(b)** Line plot of abundance at 41 hours of *C. difficile* grown in five *D. piger* spent medias in the presence of metronidazole (MTZ). The x-axis is semi-log scale. Points indicate biological replicates. Line represents mean of n=4 biological replicates. **(c)** Line plot of abundance of four clinical *C. difficile* isolates in the presence of metronidazole. Each x-axis is semi-log scale. Y-axis is integral of *C. difficile* OD600 from 0 to 48 hours. Points indicate biological replicates. Line represents mean of n=3 biological replicates. Asterisks indicate significant difference (*P < 0.05, **P < 0.01, ***P < 0.001) according to an un-paired t-test. **(d)** Line plot of abundance at 48 hours of *C. difficile* grown in a rich chemically defined media ("DM38") in the presence of metronidazole. The x-axis is semi-log scale. For the *D. piger* coculture condition, OD600 is calculated OD600 (OD600 multiplied by relative abundance from 16S sequencing). Points indicate biological replicates. Line represents mean of n=6 (fresh media) or n=3 (DP SM) biological replicates. Asterisks indicate significant difference (*P < 0.05, **P < 0.01, ***P < 0.001) according to an un-paired t-test. **(e)** Line plot of abundance at 48 hours of *C. difficile* grown in fresh media in the presence of metronidazole that was incubated in fresh media or *D. piger* spent media. The x-axis is semi-log scale. Points indicate biological replicates. Line represents mean of n=4 biological replicates. No significant difference ("ns", p>0.05) according to an un-paired t-test. **(f)** Line plot of abundance at 48 hours of *C. difficile* grown in fresh media, *D. piger* spent media, or *D. piger* spent media with fresh media spike. The x-axis is semi-log scale. Points indicate biological replicates. Line represents mean of n=4 biological replicates. Asterisks indicate significant difference (*P < 0.05, **P < 0.01, ***P < 0.001, "ns" P>0.05) according to an un-paired t-test.

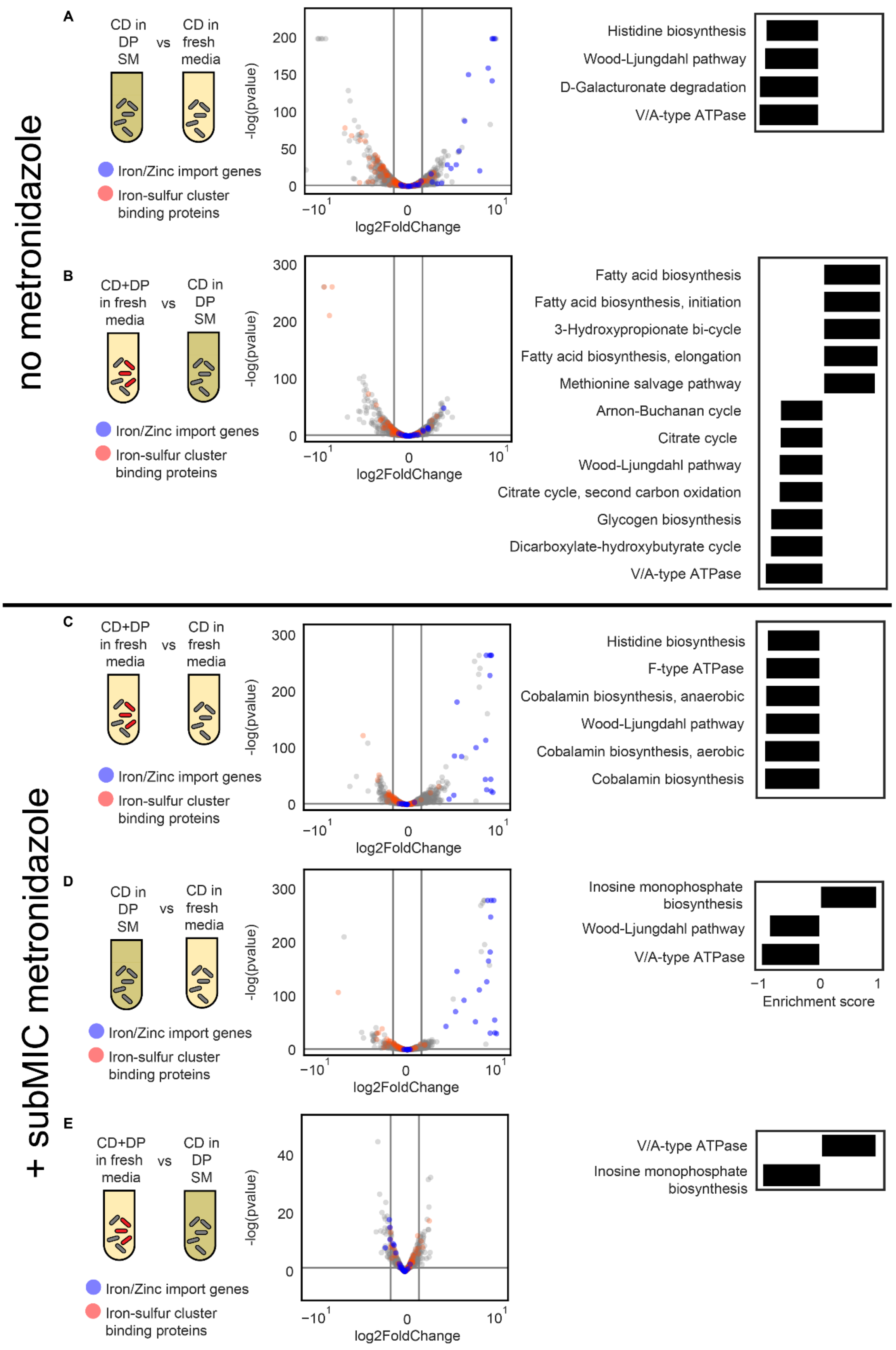

no metronidazole
+ subMIC metronidazole
A
CD in DP SM
vs
CD in fresh media
Iron/Zinc import genes
Iron-sulfur cluster binding proteins
-log(pvalue)
log2FoldChange
Histidine biosynthesis
Wood-Ljungdahl pathway
D-Galacturonate degradation
V/A-type ATPase
B
CD+DP in fresh media
vs
CD in DP SM
Fatty acid biosynthesis
Fatty acid biosynthesis, initiation
3-Hydroxypropionate bi-cycle
Fatty acid biosynthesis, elongation
Methionine salvage pathway
Arnon-Buchanan cycle
Citrate cycle
Wood-Ljungdahl pathway
Citrate cycle, second carbon oxidation
Glycogen biosynthesis
Dicarboxylate-hydroxybutyrate cycle
V/A-type ATPase
C
CD+DP in fresh media
vs
CD in fresh media
Histidine biosynthesis
F-type ATPase
Cobalamin biosynthesis, anaerobic
Wood-Ljungdahl pathway
Cobalamin biosynthesis, aerobic
Cobalamin biosynthesis
D
CD in DP SM
vs
CD in fresh media
Inosine monophosphate biosynthesis
Wood-Ljungdahl pathway
V/A-type ATPase
Enrichment score
E
CD+DP in fresh media
vs
CD in DP SM
V/A-type ATPase
Inosine monophosphate biosynthesis

**Supplementary Figure 10: Changes in expression of metabolic pathways and iron-related genes in *C. difficile.*** **(a) (b) (c) (d) (e)** Left: Schematic of two conditions being compared. Middle: Volcano plot of change in *C. difficile* gene expression. Gray vertical lines indicate 2-fold change (1 in log2) and gray horizontal line indicates statistical significance (p=0.05). Blue indicates genes annotated to be involved in iron or zinc import. Red indicates genes predicted to contain iron-sulfur clusters by MetalPredator. Right: Enriched KEGG modules in *C. difficile*. All KEGG pathways with significant enrichment scores from Gene Set Enrichment Analysis (GSEA) are shown.

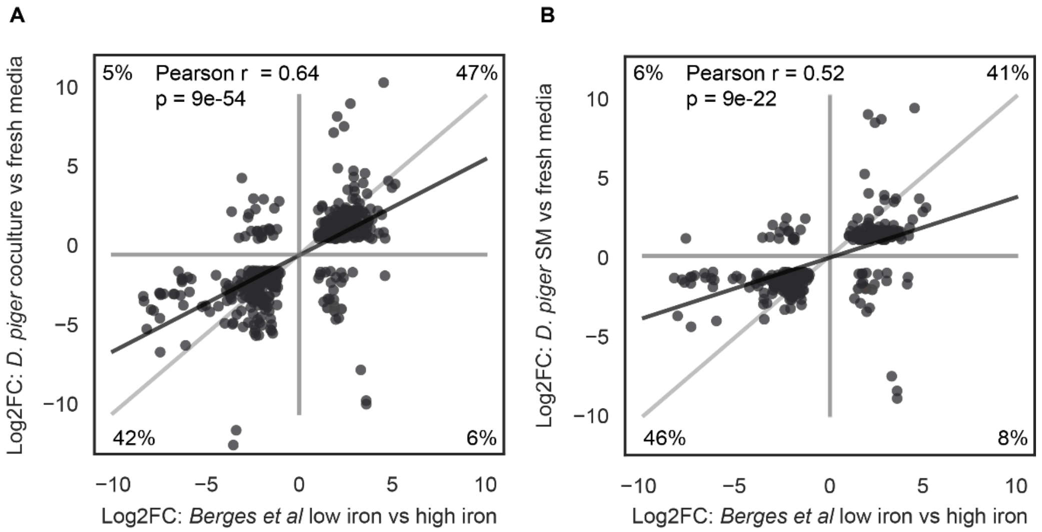


**Supplementary Figure 11: Correlation of change in gene expression between *D. piger* coculture and fresh media with change in gene expression between low iron and high iron media in Berges et al. (a) (b)** Scatterplots of comparison of gene expression in gene orthologs between CDR20291 (this study) and CD630Δerm (Berges et al). Genes only shown if differentially expressed in both studies. X-axis: log2 fold change between low iron (0.2 μM) and high iron (15 μM) media in Berges et al study. Y-axis: log2 fold change between *C. difficile* in *D. piger* coculture (panel A) or *C. difficile* in *D. piger* spent media (SM) (panel B) and *C. difficile* in fresh media in this study. Each point indicates a gene. Percentages indicate percentage of genes in each quadrant. Black line indicates best fit linear regression for all data points. Gray line is y=x.

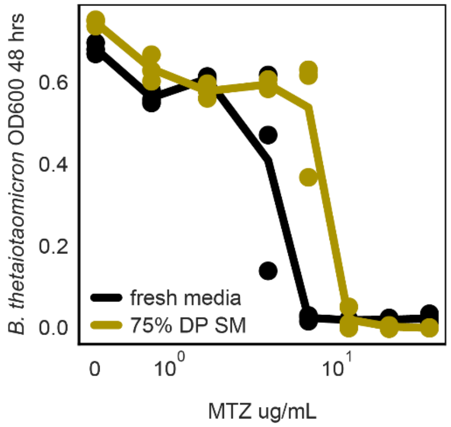


**Supplementary Figure 12: *D. piger* supernatant increases *B. thetaiotaomicron* metronidazole tolerance.** Lineplot of abundance at 48 hours of *B. thetaiotaomicron* grown in fresh media or in 75% *D.*

*piger* spent media:25% fresh media. The x-axis is semi-log scale. Points represent biological replicates. Lines indicate average of n=3 biological replicates.

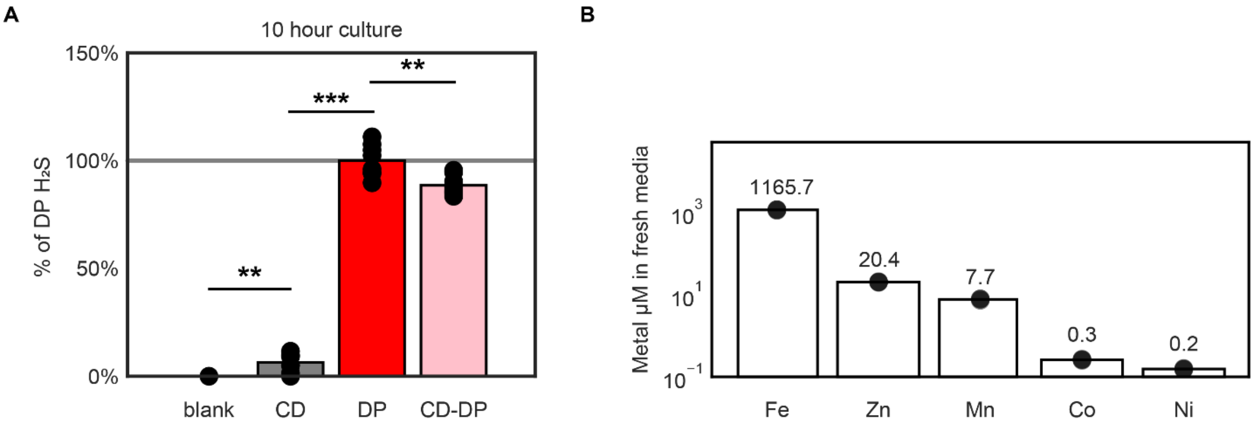


**Supplementary Figure 13: Concentrations of hydrogen sulfide and divalent transition metals. (a)** Barplot of amount of hydrogen sulfide in multiple conditions after 10 hours of incubation. "Blank" is fresh media control. Y-axis is percentage of the amount of hydrogen sulfide in *D. piger* monospecies. Points represent mean of n=2 technical replicates. Bar represents mean of n=8 biological replicates. Asterisks indicate significant difference (*$P < 0.05$, **$P < 0.01$, ***$P < 0.001$, "ns" $P>0.05$) according to an un-paired t-test. **(b)** Barplot of concentration of divalent transition metals in fresh media. Points represent mean of n=2 technical replicates. Bar represents mean of n=3 biological replicates.

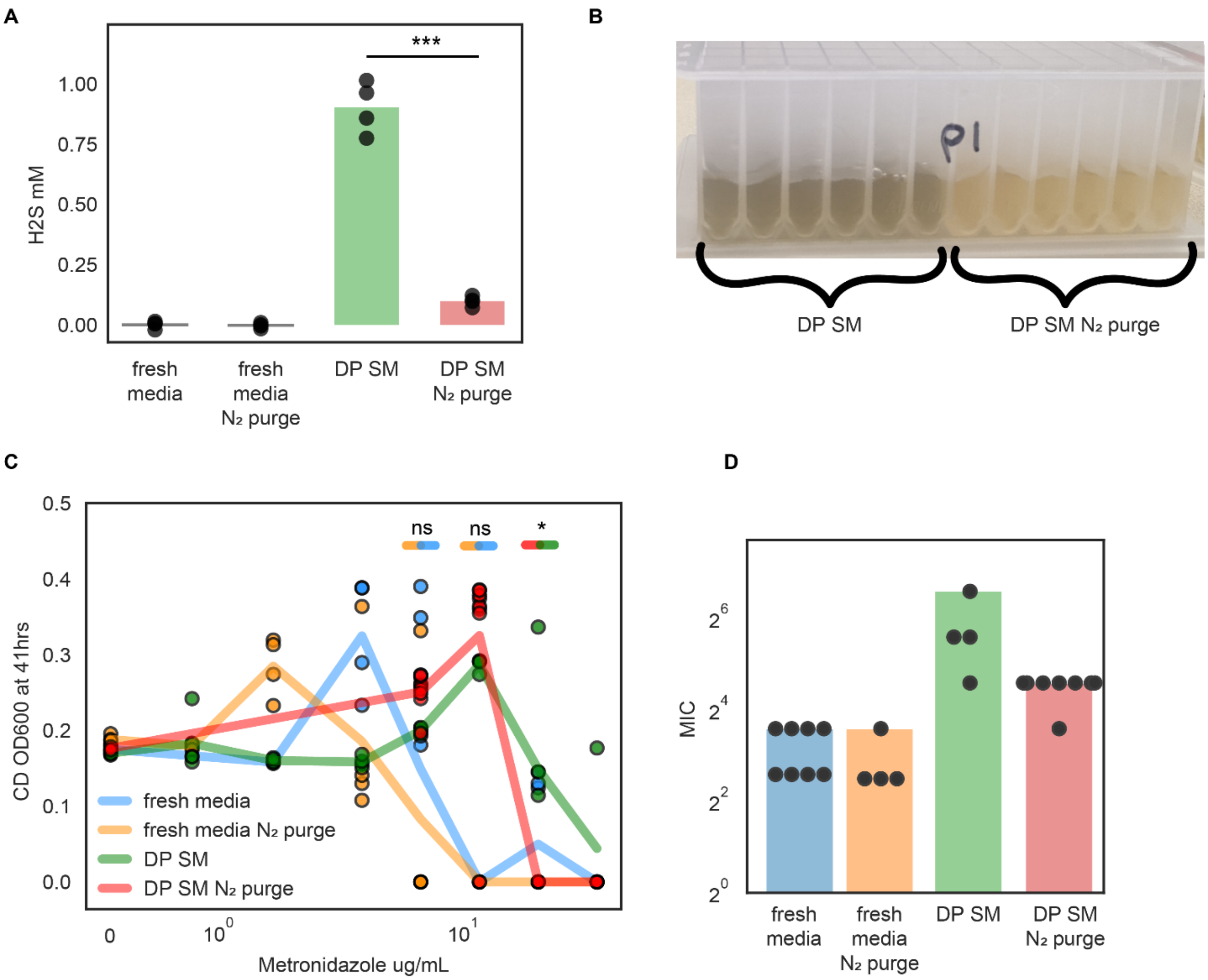


**Supplementary Figure 14: Nitrogen purge of *D. piger* spent media decreases hydrogen sulfide concentration, metal precipitate, and *C. difficile* metronidazole MIC. (a)** Barplot of hydrogen sulfide concentration of medias. Points represent n=4 biological replicates. Bar represents average of biological replicates. **(b)** Image of *D. piger* spent media 2 hours after nitrogen purge (right) or no treatment (left). Precipitation of black ferrous sulfide turns media a darker color. **(c)** Lineplot of OD600 at 41 hours of *C. difficile* grown in different medias in the presence of metronidazole. The x-axis is semi-log scale. Points represent biological replicates. Lines indicate average of n=4 to n=8 biological replicates. **(d)** Barplot displaying MIC of data shown in C. Points represent MIC of n=4 to n=8 biological replicates. Bar represents MIC determined based on average OD600 of n=4 to n=8 biological replicates.

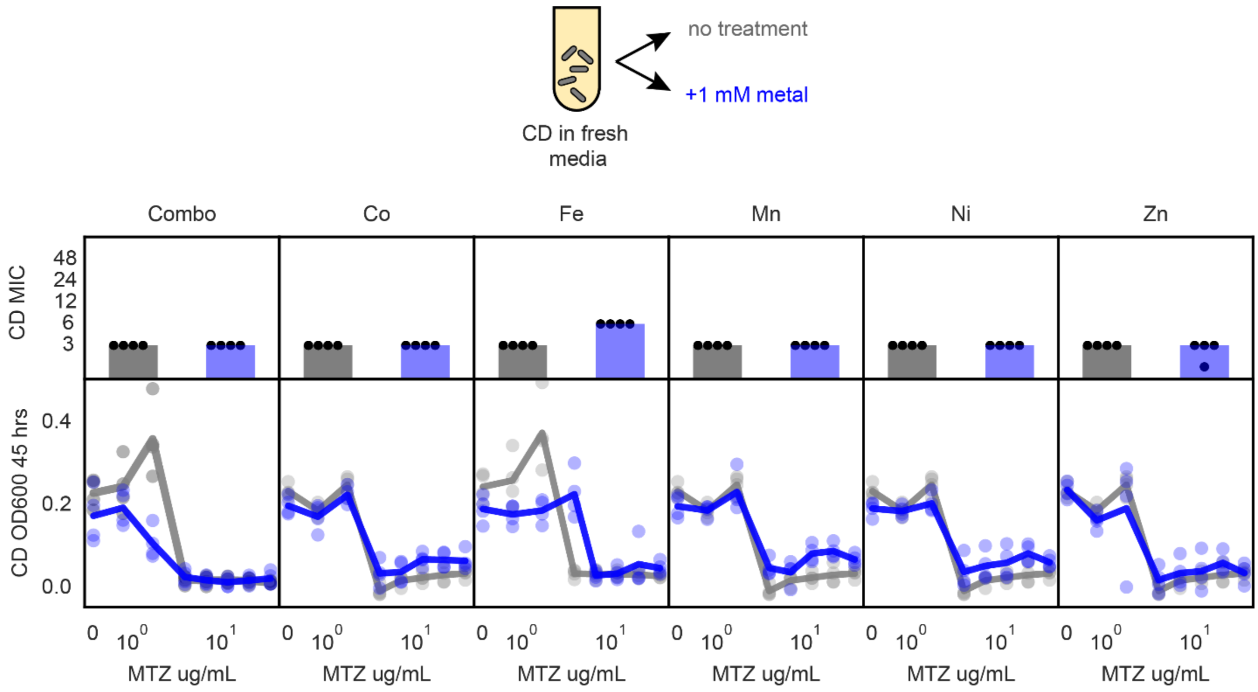


**Supplementary Figure 15: Supplementation of fresh media with metals does not decrease *C. difficile* tolerance to metronidazole**. Lineplots and barplots of *C. difficile* metronidazole susceptibility in fresh media with and without metal supplementation. (Top) Barplots of *C. difficile* metronidazole MIC in untreated fresh media (gray) and media with 1mM metal supplementation (blue). In metal combination condition ("Combo") all five metals were supplemented at 1 mM. Points represent MIC of n=4 biological replicates. Bar represents MIC determined based on average OD600 of n=4 biological replicates. (Bottom) Lineplots of OD600 at 45 hours of *C. difficile* grown in untreated fresh media (gray) or media supplemented with 1 mM metal (blue) in the presence of metronidazole (MTZ). Each x-axis is semi-log scale. Points represent biological replicates. Lines indicate average of n=4 biological replicates.

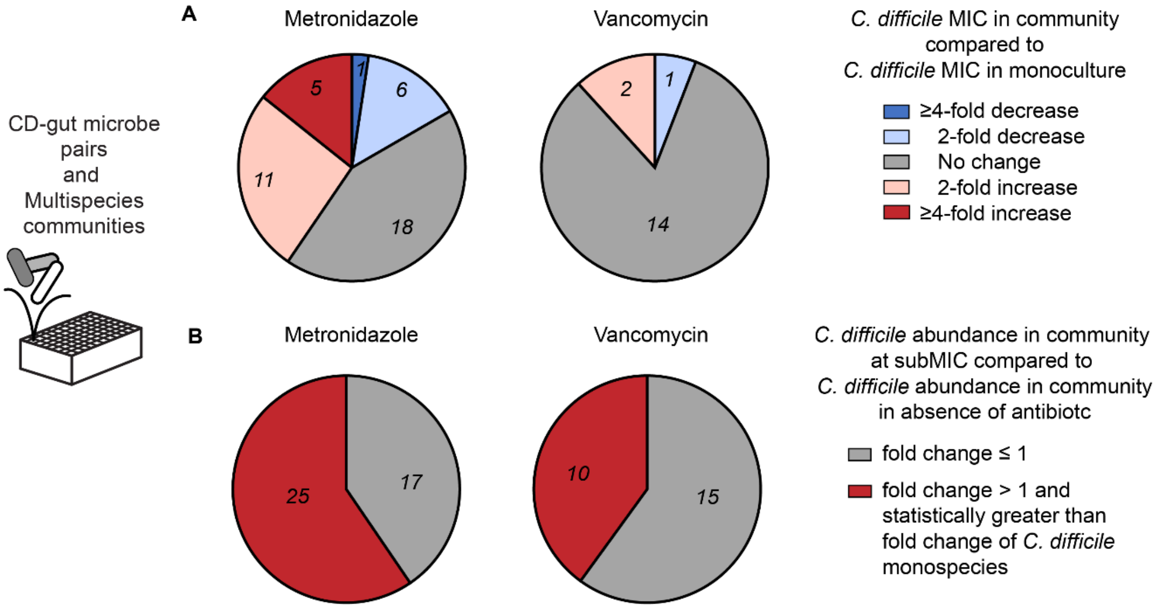


**Supplementary Figure 16: Summary of alteration of C. difficile antibiotic response in both pairwise and multispecies communities.** **(a)** Pie charts of change in *C. difficile* MIC in communities compared to monospecies for pairwise and multispecies community data. **(b)** Pie charts of growth enhancement of *C. difficile* in pairwise and multispecies communities. Growth enhancement is defined as a maximum subMIC fold change that is greater than one and is greater than the maximum subMIC fold change in *C. difficile* monospecies for that antibiotic (i.e., greater than 1.61 for metronidazole).
